# Organization and composition of apicomplexan kinetochores reveal plasticity in chromosome segregation across parasite modes of division

**DOI:** 10.1101/2021.11.03.466924

**Authors:** Lorenzo Brusini, Nicolas Dos Santos Pacheco, Dominique Soldati-Favre, Mathieu Brochet

## Abstract

Kinetochores are multiprotein assemblies directing mitotic spindle attachment and chromosome segregation. In apicomplexan parasites, most known kinetochore components and associated regulators are apparently missing, suggesting a minimal structure with limited control over chromosome segregation. In this study, we use interactomics combined with deep homology searches to identify six divergent eukaryotic components, in addition to a set of eight apicomplexan kinetochore proteins (AKiTs) that bear no detectable sequence similarity to known proteins. The nanoscale organization of the apicomplexan kinetochore includes four subdomains, each displaying different evolutionary rates across the phylum. Functional analyses confirm AKiTs are essential for mitosis and reveal architectures parallel to biorientation at metaphase. Furthermore, we identify a homolog of MAD1 at the apicomplexan kinetochore, suggesting conserved spindle assembly checkpoint signaling. Finally, we show unexpected plasticity in kinetochore composition and segregation throughout the parasite lifecycle, indicating diverse requirements to maintain fidelity of chromosome segregation across apicomplexan modes of division.

## Introduction

Eukaryotic chromosome segregation occurs along a spindle formed of microtubules. In order to segregate DNA into daughter cells, the spindle must interact with chromosomes. Most organisms achieve specificity in this interaction by binding to chromosomal sites called centromeres, distinguished by the histone H3 variant centromere protein A (CENP-A) (Fukagawa & Earnshaw, 2014; Westhorpe & Straight, 2013). Onto the centromere assembles the kinetochore, a hierarchical assembly and molecular machine that links chromosomes to the spindle (Cheeseman & Desai, 2008). The kinetochore is formed of multiple compartments, each composed of different protein complexes. In animals and fungi, the constitutive centromere-associated network (CCAN) forms the inner kinetochore (Foltz et al., 2006), whilst at the onset of mitosis the outer kinetochore KMN network is recruited, formed of KNL1, MIS12 and NDC80/NUF2 complexes (Cheeseman et al., 2006). It is the KMN network that serves as both the microtubule-binding component of the kinetochore and a landing pad for either SKA or DASH complexes that strengthen microtubule attachment in animals and fungi (Helgeson et al., 2018; Lampert et al., 2010, 2013), respectively, in addition to the Spindle Assembly Checkpoint (SAC), a surveillance system that ensures faithful chromosome segregation (Varma & Salmon, 2012). Fidelity requires that sister chromatids are bi-oriented at metaphase, kinetochores bound by microtubules emanating from opposing spindle poles (Lampson & Cheeseman, 2011). Bi-orientation is achieved through an error-correction process mediated by a “wait” signal from the Mitotic Checkpoint Complex (MCC) (Musacchio & Salmon, 2007). This diffusable inhibitor prevents precocious activation of the anaphase-promoting complex/cyclosome (APC/C). Upon bi-orientation, the APC/C is liberated and initiates the cascade that culminates in cleavage of cohesin and segregation of sister chromatids into daughter cells.

The conservation of kinetochore proteins varies greatly across organisms (Tromer et al., 2019; van Hooff et al., 2017). Whilst components of the KMN network have been detected in all studied eukaryotes (D’Archivio & Wickstead, 2017), most components of the CCAN are not readily identified (Przewloka et al., 2007; Westermann & Schleiffer, 2013). In particular, members of the phylum Apicomplexa lack almost all clearly identifiable components of the CCAN and SAC as described in animals and fungi, in addition to the majority of the KMN network (D’Archivio & Wickstead, 2017; van Hooff et al., 2017). This phylum groups a large number of obligate intracellular parasites of considerable medical and veterinary relevance, including the malaria parasite *Plasmodium* and *Toxoplasma*, causative agent for toxoplasmosis. In addition to a widespread lack of “canonical” kinetochore components, apicomplexan parasites appear to divide quite differently to most of the cells of their hosts (Francia & Striepen, 2014; Gerald et al., 2011; Striepen et al., 2007). Flexibility in the scale of amplification and modes of division, often adapting to the size and environment of different host cells and tissues, suggests division checkpoints in these parasites may be very different to those described in animals (Alvarez & Suvorova, 2017; D. E. Arnot & Gull, 1998). Asexual forms of *T. gondii* divide in tissues of the mammalian intermediate host by a mechanism termed endodyogeny, producing two daughter cells from a single round of fission within the mother cell, which is consumed as the offspring mature. Uncoupling nuclear divisions from the cell cycle allows certain apicomplexan parasites to successively replicate their genome in the absence of cytokinesis. In red blood cells, *Plasmodium* species divide by schizogony to produce multi-nucleated coenocytes. Similar division modes occur in liver hepatocytes and beneath the mosquito midgut basal lamina (Aly et al., 2009). Karyokinesis is asynchronous, resulting in non-geometric expansion. Following the last round of division, daughter cells either bud internally or from the surface of the mother plasmalemma (David E Arnot et al., 2011; Speer & Dubey, 2001). During host-to-vector transmission, a mosquito blood meal induces capacitated male gametocytes (microgametocytes) to produce a polyploid nucleus that then divides to form 8 haploid gametes. Microgametes fertilize female gametes (macrogametes), shortly followed by meiosis and differentiation producing characteristic banana-shaped ookinetes.

Three organelles contain DNA in apicomplexan parasites. In addition to the mitochondrion, the nucleus and a remnant of secondary symbiosis called the apicoplast each have their own distinct segregation cycle. The nucleus undergoes a largely closed mitosis, an intranuclear spindle nucleates from spindle poles and links centromeres to kinetochore-like structures maintained close to the nuclear periphery (David E Arnot et al., 2011; Dubremetz, 1973; Farrell & Gubbels, 2014; Zeeshan et al., 2020). Current evidence suggests apicomplexan parasites segregate their chromosomes with fidelity (Iwanaga et al., 2010, 2012). However, in light of the apparent absence of most kinetochore and checkpoint proteins, how these organisms maintain faithful chromosome segregation remains unknown.

Here, we identify the repertoires of four biochemically stable compartments of apicomplexan kinetochores that include at least six divergent eukaryotic components, in addition to a set of eight for which we did not detect similarity with known proteins. Despite sequence disparity, these apicomplexan kinetochore proteins (AKiTs) display modes of chromosome segregation parallel to metaphase to anaphase transition and include a homolog of the spindle assembly checkpoint protein MAD1, that we show also localizes to apicomplexan kinetochores. Furthermore, we show plasticity in kinetochore composition and segregation between lifecycle stages, suggestive of unique requirements and regulation between modes of division. Finally, AKiTs are required for proper parasite proliferation, identifying the apicomplexan kinetochore as an excellent therapeutic candidate for selective inhibition.

## Results

### NUF2 and SKA2 assemble at the nuclear periphery in the malaria parasite

Whilst the majority of kinetochore components are not clearly detectable in apicomplexan parasites (van Hooff et al., 2017), centromeric proteins (CENPs) A, C and E, the SKA component SKA2 and the NDC80/NUF2 complex are present (Fig. S1A), and NDC80/HEC1 has recently been described as a *bona fide* kinetochore marker in the rodent malarial parasite *Plasmodium berghei* (Zeeshan et al., 2020). To understand whether apicomplexan kinetochore proteins bear similarities to their animal and fungal cousins, we took a pan-localization approach by tagging *P. berghei* components related to different animal kinetochore sub-complexes. Candidate proteins were localized throughout asexual blood-stages and during sexual development that occurs upon host-to-mosquito transmission. Immunoblotting confirmed expression of fusion proteins with the expected mobility (Fig. S1B & C) corresponding to the integration of the coding sequence for mScarlet-I (mSc) or mNeonGreen fused with triple hemagglutinin- epitope tag (mNG-3xHA) at the C-terminus of endogenous NUF2 (PBANKA_0414300) and SKA2 (PBANKA_0405800), respectively.

In agreement with previously localized *Plasmodium* centromeres (Perrin et al., 2021; Verma & Surolia, 2014) and kinetochores (Zeeshan et al., 2020), location and movement of NUF2 and SKA2 fusion proteins was restricted to the nuclear periphery during progression of asexual blood-stage divisions (Fig. 1A). Protein levels were undetectable in G1 phase in intraerythrocytic ring stages and first seen in the nucleus at the onset of DNA replication during trophozoite development, then accumulating as punctate foci concomitant with chromosome segregation and formation of daughter nuclei. Signal for both NUF2 and SKA2 fusions reduced to below detectability in fully budded schizonts.

**Figure 1.**
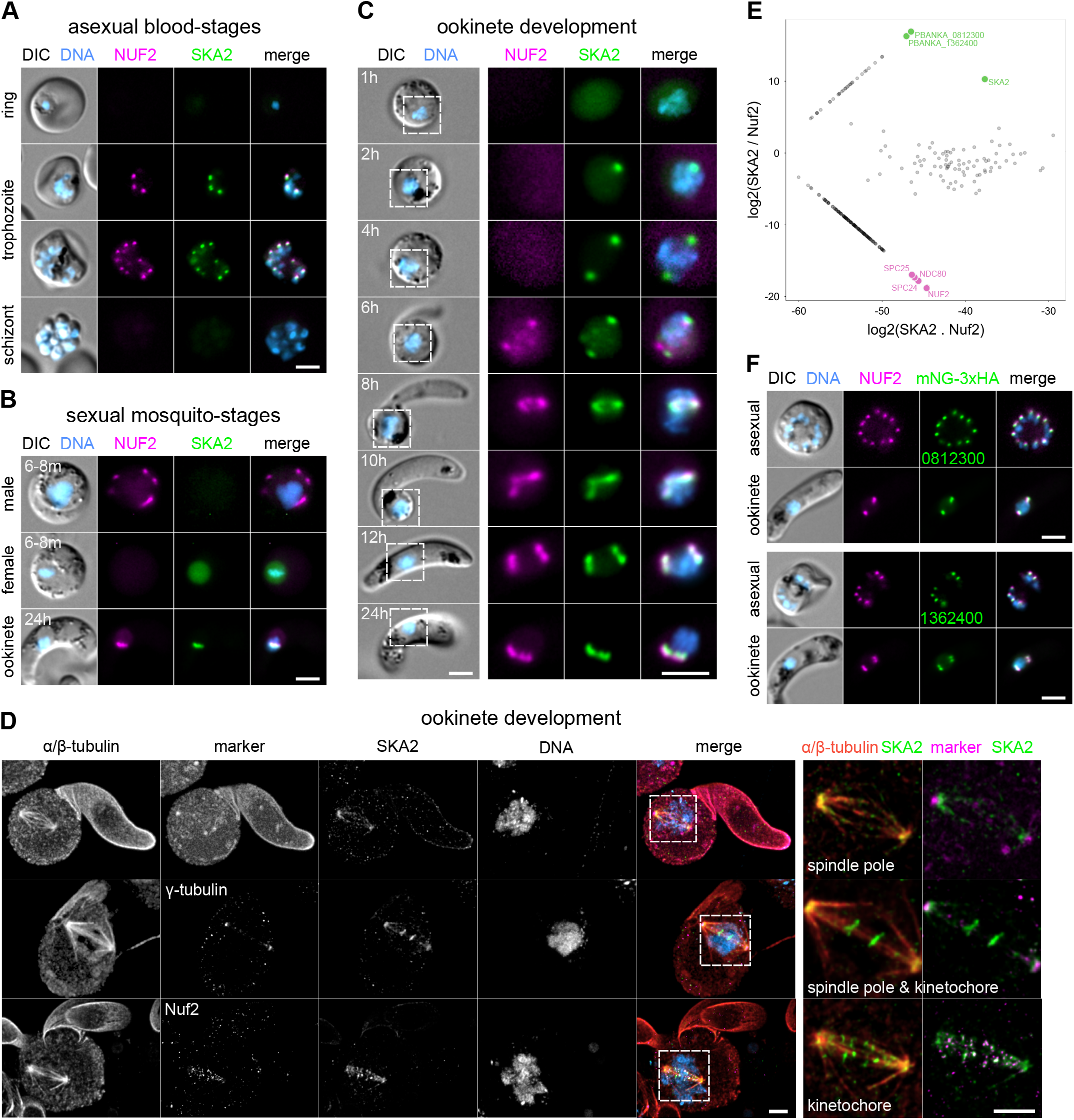
*Plasmodium* species encode a biochemically stable SKA complex. Micrographs of live native fluorescence in malaria parasites expressing tagged kinetochore components NUF2-mScarlet-I (magenta) and SKA2-mNeonGreen fused to triple hemagglutinin-epitope tag (green) during asexual blood-stage proliferation (A), sexual mosquito-stages (B), and throughout meiosis and differentiation in developing ookinetes (C). Counter-staining of DNA with Hoechst 33342 (cyan) and differential interference contrast images (DIC) are also shown. Bar: 2 μm. (D) Ultrastructure Expansion Microscopy revealed that tagged-SKA2 localizes to spindle poles (identified by γ-tubulin and pan-proteomic NHS-ester counter-stainings), along the spindle (α/β-tubulin) and at kinetochores (NUF2) during meiosis. Counter-staining of DNA with Hoechst 33342 (cyan) also shown. Bar: 1 μm. (E) Label-free semiquantitative mass spectrometry showing relative enrichment of proteins identified following immunoprecipitations of tagged kinetochore components NUF2-3xHA (magenta) and SKA2-3xHA (green). Signals from the NDC80 complex and two novel SKA2-interacting proteins are highlighted. Intensities of proteins not detected for a specific immunoprecipitation are set to an arbitrary minimum value. (F) Live native fluorescence validated SKA2-interacting proteins tagged with mNG-3xHA localize to kinetochore foci during asexual blood stages and ookinete development.

Unexpectedly, SKA2 and NUF2 fusion proteins showed very different localizations during *Plasmodium* sexual development (Fig. 1B). Whilst “rod-like bridges”, as previously described for components of the NDC80/NUF2 complex (Zeeshan et al., 2020), were visible for NUF2 during the three rounds of DNA replication and mitosis that occur at microgametogenesis – SKA2 signal at kinetochore foci was not clearly detectable. A similar dichotomy was seen post-activation of the macrogametocyte, NUF2 spreading sparingly across the cytoplasm compared to SKA2 residing primarily in the nucleus. However, 24-hours post-fertilization and meiosis, both fusion proteins united as four distinct nuclear foci in fully developed banana-shaped ookinetes. Time-course fluorescence microscopy further revealed a hierarchical assembly for SKA2 and NUF2 throughout ookinete development (Fig. 1C). SKA2 first accumulated as a single focus at the nuclear periphery between 1 - 2 hours post-fertilization, which duplicated and migrated to opposing nuclear poles between 2 - 4 hours. On the other hand, NUF2 foci were first seen between 4-6 hours of development, at which point SKA2 signal stretched along a spindle-like structure connecting opposing poles. Signal from both fusion proteins stretched along the central spindle by 8 - 10 hours. Two successive rounds of asynchronous duplication and migration ultimately formed four puncta at the nuclear periphery by 12 hours of development.

Given the limitations in defining nanometer-scale structures by conventional fluorescence microscopy, we applied Ultrastructure Expansion Microscopy (U-ExM) during ookinete development to increase resolution (Bertiaux et al., 2021; Gambarotto et al., 2019). Pan-proteomic (NHS-ester), subpellicular/spindle microtubule and nuclear stainings identified parasite shape and ultrastructure were well preserved following expansion of between 4.5 - 4.6x, and, alongside the localizations of tagged SKA2 and NUF2, resolved kinetochores along the *Plasmodium* meiotic spindle (Fig. 1D). SKA2 accumulated primarily at three locations during meiosis in ookinetes. Consistent with our live fluorescence microscopy, the majority of SKA2 localized to spindle poles, closely associated to γ-tubulin. However, upon formation of a diamond shaped bipolar spindle, additional foci were obvious both along the spindle and at the spindle equator, co-localizing with NUF2 at kinetochores.

These observations indicate that SKA2 localizes to both the spindle and kinetochores in the malaria parasite, similarly to SKA components in animals (Hanisch et al., 2006).

### NUF2 and SKA2 form distinct biochemically stable complexes in Plasmodium

In models, the NDC80 complex is formed of two heterodimers: NDC80 with NUF2, and SPC24 with SPC25. The metazoan SKA complex forms an independent three component complex: SKA1 - 3 (Hanisch et al., 2006; Welburn et al., 2009). In *P. berghei*, a divergent SPC24 ortholog was recently identified as the final missing component of the NDC80/NUF2 complex, copurifying with NDC80 following affinity purification from chemically cross-linked cells (Zeeshan et al., 2020). To identify interacting proteins for NUF2 and SKA2, we used affinity purification of 3xHA tagged proteins from synchronized mitotic gametocytes under non cross-linking conditions and combined label-free semiquantitative mass spectrometry (Trudgian et al., 2011) to estimate enrichment of interacting proteins. We compared samples by integrated spectral intensities (Fig. 1E), reasoning that if NDC80 and SKA2 assemble independent kinetochore complexes in *Plasmodium*, stable interactors would be evident in each pull down relative to one another. Supporting the notion of a biochemically stable NDC80/NUF2 complex, NDC80, SPC25 and recently described SPC24 were the most abundant proteins co-purifying with NUF2 relative to SKA2. Similarly, the abundance of two hypothetical proteins of unknown function (Gene IDs: PBANKA_0812300 and PBANKA_1362400) strongly suggests the presence of a three SKA component complex in the malaria parasite. To validate our biochemical approach, we tagged each SKA2 interacting protein with mNG-3xHA, alongside the marker for the outer kinetochore NUF2-mSc (Supp. Fig. S1D & E). Both tagged PBANKA_0812300 and PBANKA_1362400 proteins showed localization patterns highly similar to that of SKA2 in the malaria parasite (Fig. 1F & S1F & G), colocalizing with NUF2 during asexual blood stage divisions, not detected during microgametogenesis and present as four nuclear foci in fully developed ookinetes.

### SKA components are required for timely kinetochore segregation in Toxoplasma

The SKA complex has been identified in all 5 - 6 eukaryotic supergroups (He et al., 2014; van Hooff et al., 2017). Despite this broad distribution, the complex is not ubiquitously detected in functionally characterized kinetochores, such as in fungi and kinetoplastids. Missing individual SKA components occurs more sporadically, suggesting spurious absences may result from lack of detection due to sequence divergence rather than genuine gene loss. Consistent with the notion of rapidly changing kinetochore protein sequences in apicomplexan parasites, BlastP of PBANKA_0812300 and PBANKA_1362400 in the “closely related” apicomplexan parasite, *Toxoplasma gondii,* yielded no significant hits (e-value ≥ 0.1). In an attempt to more sensitively identify SKA2-interacting proteins, we employed a similar iterative Hidden Markov Model (HMM) profiling strategy to as described (Koreny et al., 2021). Briefly, HMMs constructed from clear homologs of PBANKA_0812300 and PBANKA_1362400 were used as search queries across an in-house database of HMMs generated from homologous groups of alveolate protein sequences (Table S1), however with the addition of manually curated HMMs generated from previously classified eukaryotic kinetochore sequences (van Hooff et al., 2017). Significant hits were concatenated iteratively to generate pan-apicomplexan-kinetochore HMMs (Supplementary HMM files). In contrast to BlastP, reciprocal HMM profile-profile comparisons identified significant similarity in both SKA2-interacting proteins, with pan-SKA HMMs as highest scoring hits (Fig. S2A & B). In particular, we reunited PBANKA_0812300 with SKA1 and PBANKA_1362400 with SKA3, and detected corresponding putative homologs in *T. gondii* TGME49_264960 and TGME49_289790, respectively.

To validate our bioinformatic approach and investigate the conservation of SKA proteins across the Apicomplexa, TgSKA1 and TgSKA2 were fused at the endogenous locus with a mini auxin-induced degron (mAID) including triple hemagglutinin (3xHA)-epitope tag, to localize and deplete protein upon addition of auxin (IAA) (Fig. S2F & G). Supporting identification of divergent apicomplexan SKA components, tagged TgSKA1 and TgSKA2 showed clear kinetochore localization during *T. gondii* tachyzoite division (Fig. 2A). Co-staining DNA and the mitotic spindle revealed SKA proteins were undetectable in G1 phase, but accumulated centrally upon a bipolar spindle during mitosis. Foci segregated at anaphase into budding daughter cells, returning to below detectability at cytokinesis. Co-expression with the kinetochore component NUF2 fused to a double Ty epitope tag (NUF2-2xTy), in addition to co-staining with antibodies directed to various intracellular organelles, revealed *T. gondii* SKA proteins are substantially displaced distal to the apicoplast and centrosome compared to kinetochores (Fig. S2E).

**Figure 2.**
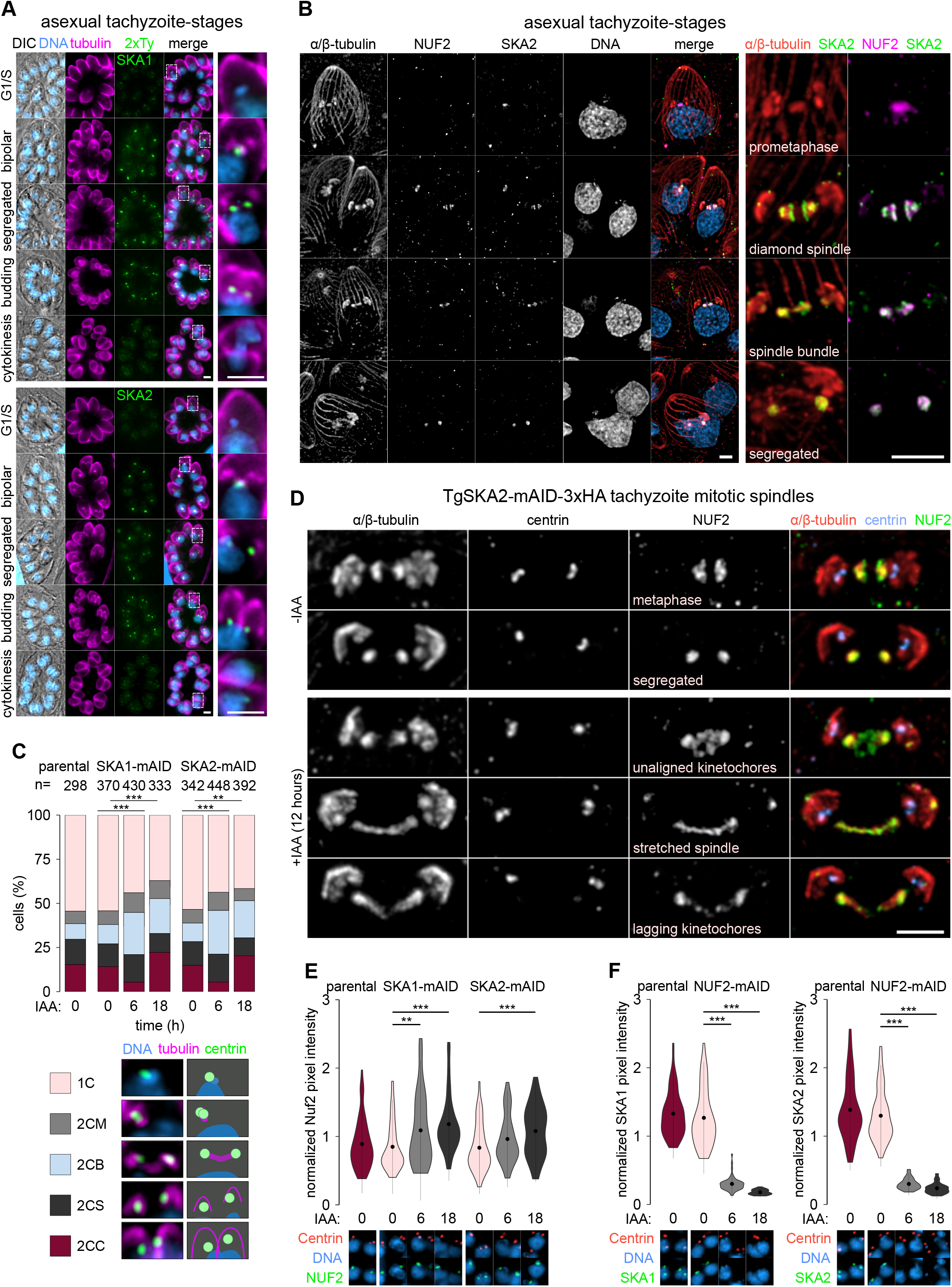
SKA components are required for kinetochore segregation in *Toxoplasma.* (A) Micrographs of fixed immunofluorescence in *Toxoplasma gondii* tachyzoites expressing kinetochore components SKA1 and SKA2 fused to a double Ty epitope (2xTy) throughout intracellular divisions. Counter-staining of DNA with DAPI (cyan), tubulin (magenta) and DIC images are also shown. Bar: 5 μm. (B) U-ExM revealed SKA2 is recruited to the tachyzoite mitotic spindle primarily at metaphase, to spindle poles (identified by α/β-tubulin counter-stain) and kinetochores (NUF2). Counter- staining of DNA with DAPI (cyan) also shown. Bar: 2 μm. (C) Morphological analysis of cells depleted for SKA1 or SKA2 tagged with mAID-3xHA shows a buildup of cells with duplicated centrosomes and bipolar spindles (2CB) compared to monopolar spindles (2CM), segregated centrosomes into daughter cell caps (2CS) and cells in cytokinesis (2CC). Assessed by immunofluorescence against *Toxoplasma* Centrin1 (green) and tubulin (magenta) (**, P < 0.01; ***, P < 0.001; Chi^2^ test). (D) U-ExM further characterized phenotypic changes post-depletion, revealing tachyzoites with stretched mitotic spindles and unaligned and lagging kinetochores during mitosis. Bar: 1 µm. Levels and localization of NUF2 (E) and SKA components (F) in cells at 0, 6 and 18 hours post-depletion of SKA and NUF2 components, respectively, assessed by immunofluorescence against Centrin1 and 2xTy tagged protein. Representative images shown below. DNA stain DAPI.

U-ExM further resolved TgSKA2 at the tachyzoite mitotic spindle (Fig. 2B). In tachyzoites, the spindle nucleates from a region closely associated to a centriolar microtubule organizing center. A short prometaphase spindle elongates to form a bipolar opposing diamond at metaphase, prior to collapse into a tight central bundle and segregation into budding daughter caps. In contrast to NUF2, which localized to the spindle at prometaphase, SKA2 signal was only detected upon diamond-spindle formation. As in the malaria parasite, the majority of SKA2 signal localized to the spindle poles, along the spindle and at the midzone.

Depletion of either TgSKA1 or TgSKA2 by IAA treatment led to a rapid and severe reduction of cell growth (Fig. S2F). Consistent with a role in nuclear chromosome segregation, vacuoles contained accumulations of DNA with no associated daughter cell body 18 hours post-depletion (Fig. S2G). This defect was evident within the first round of mitosis 6 hours post-depletion, with an accumulation of parasites with monopolar and bipolar spindles and budding daughter caps (Fig. 2C). U-ExM revealed cells stalled during mitosis, with elongated mitotic spindles and misaligned and lagging kinetochores, not seen in non-induced and normal cells (Fig. 2D).

To investigate the hierarchy of outer kinetochore components in *T. gondii*, we looked at the recruitment and co-dependency of the NDC80/NUF2 complex component NUF2 relative to SKA1 or SKA2, in cells depleted for either protein (Fig. 2E & F). Whilst neither SKA component was required for recruitment of NUF2 onto kinetochore foci (instead a surprising increase of NUF2 signal was seen post-SKA depletion), kinetochore assembly of SKA1 and SKA2 was abolished in mitotic tachyzoites depleted for NUF2.

Taken together, apicomplexan kinetochores require SKA components for proper mitosis and furthermore suggest that maintenance of the SKA complex at kinetochores is dependent on, and possibly downstream to, assembly of the NDC80/NUF2 complex.

### Apicomplexan Kinetochore protein 1 (AKiT1) is a component of the Plasmodium kinetochore

The apparent absence of CCAN components outside of CENP-C, in addition to Mis12/KNL1 of the KMN network, questions what bridges the outer kinetochore to the centromere in apicomplexan parasites? NDC80 and SKA complexes appear to be biochemically distinct sets and affinity purification of NUF2 and SKA2 from *P. berghei* gametocytes did not clearly enrich for additional candidates. To investigate the possibility of less stable inter-complex associations, we employed a proximity-based approach of affinity purification under conditions of formaldehyde cross-linking and compared spectral intensities to controls without cross-linking (Fig. 3A). In addition to components of the minichromosome maintenance (MCM) complex; MCM3 and MCM6, two proteins were most enriched upon crosslinking (green). One of these proteins is a homolog of STU2 (PBANKA_1337500; Fig. S3A), a protein required for the correction of kinetochore-spindle attachment errors and bipolar spindle stability during chromosome bi-orientation (Miller et al., 2016, 2019). In contrast, a hypothetical protein of unknown function (PBANKA_0621300) bore no clear sequence similarity to known animal or fungal proteins, whilst orthologs were identified across apicomplexan genomes (Fig. S3B). In agreement with biochemical interaction, the tagged protein showed clear co-localization with NUF2 throughout asexual blood-stage divisions (Fig. 3B). However, in contrast to previously localized SKA components, foci were also seen across spindle bundles during microgametogenesis (Fig. 3C & S3C). Furthermore, the identified protein accumulated as foci in the nuclei of female gametes prior to fertilization, suggesting kinetochore recruitment prior to outer kinetochore NDC80/NUF2 complex assembly. Given biochemical affinities, kinetochore localization and conservation across apicomplexan organisms, we named the protein Apicomplexan Kinetochore protein 1 (AKiT1).

**Figure 3.**
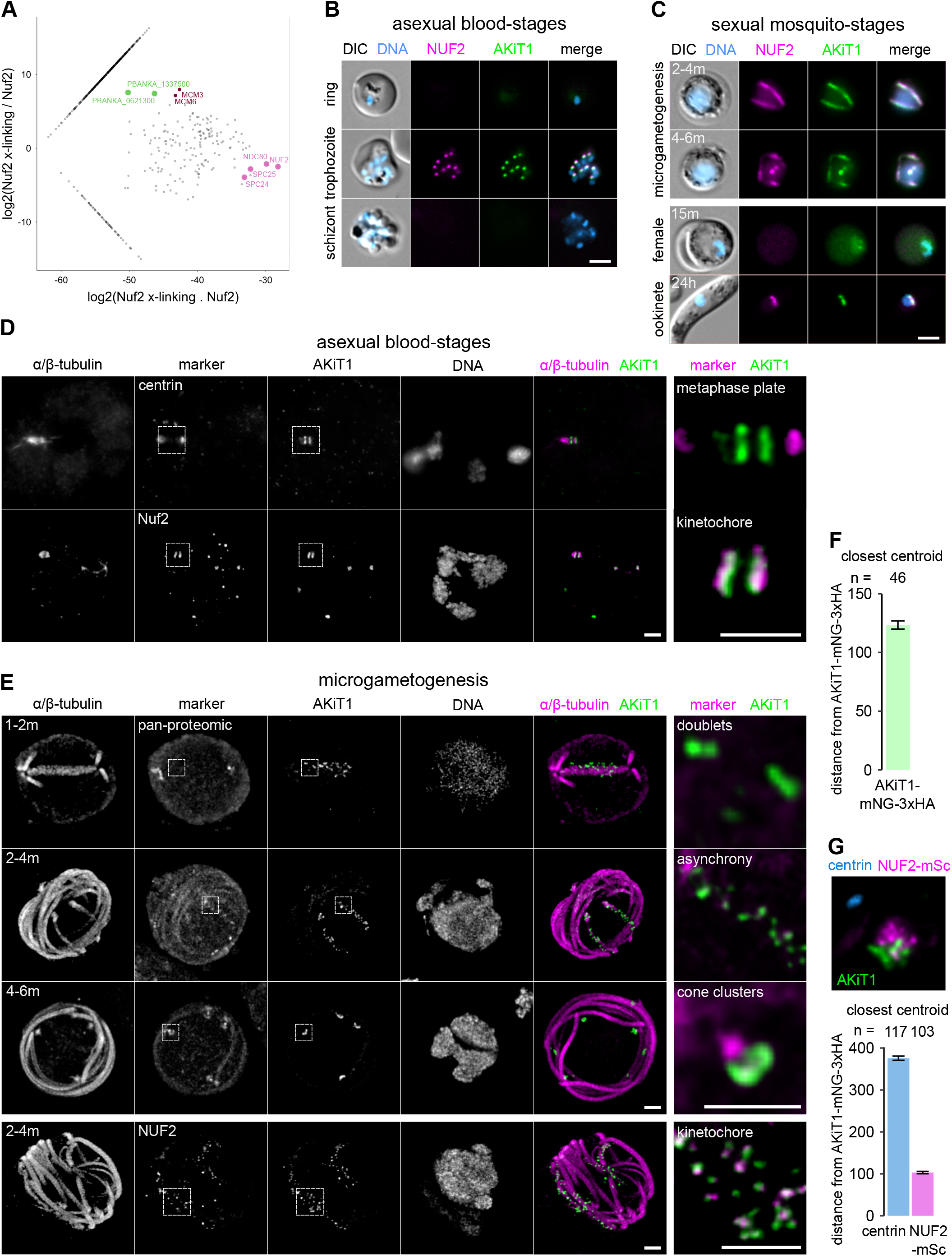
AKiT1 belongs to a newly identified compartment of the *Plasmodium* kinetochore. (A) Relative enrichment of proteins following immunoprecipitation of NUF2-3xHA under conditions of formaldehyde cross-linking compared to non cross-linked cells. Signals from NDC80 complex components and most enriched proteins are highlighted. Live fluorescence in malaria parasites expressing tagged kinetochore components NUF2-mScarlet-I (magenta) and AKiT1-mNeonGreen-3xHA (green) during asexual blood-stage proliferation (B) and microgametogenesis (C). Bar: 2 μm. U-ExM revealed alignment of AKiT1 foci at kinetochores (identified by NUF2 counter-stain) during asexual blood-stage divisions (D), whilst dispersed along the mitotic spindle (tubulin) prior to segregation during microgametogenesis (E). DNA, centrin and pan-proteomic stainings are also shown. Bar: 1 μm. Distance between closest centroids of AKiT1 along the microgametocyte spindle (F), and of NUF2 and centrin relative to AKiT1 (G). Numbers indicate total number of foci. Bars show S.E.M.

U-ExM of PbAKiT1 alongside spindle pole and kinetochore markers in asexual blood-stage cells revealed that centrosome migration and spindle elongation were accompanied by alignment of kinetochores at the spindle midzone (Fig. 3D), forming a structure reminiscent of the metaphase plate as described in animal cells. Interestingly, spindle length changed drastically between asexual and sexual-stage cells (Fig. 3D - E & S3D). During the first round of microgametocyte mitosis, PbAKiT1 localized along the spindle as pairs of kinetochore foci (Fig. 3F; mean distance apart 139 nm ± 4 nm). Kinetochores then fired asynchronously to spindle poles (Fig. 3F & S3D) – which may go some way to explaining the “rod-bridge” phenotype described in cells imaged at lower resolutions (Zeeshan et al., 2020; and this study). Two additional rounds of mitosis ultimately produced kinetochore foci arranged into 6 - 8 cone-shaped clusters at the nuclear periphery. Crucially, additional staining of NUF2 identified sub-kinetochore localizations and a clear bipartite architecture in segregated kinetochore clusters, with the distinction of PbAKiT1 at the inner kinetochore relative to PbNUF2 at the outer kinetochore and centrin at the centrosome (Fig. 3G; mean distances of 103 nm ± 3 nm and 422 nm ± 7 nm, respectively).

### AKiT1 is essential for kinetochore segregation and spindle function in Toxoplasma

Previous genome-wide screenings have indicated that AKiT1 is essential for blood-stage proliferation in both human and rodent malaria parasites (Zhang et al., 2018). Furthermore, despite several attempts, we were unable to interrogate AKiT1 function using conditional approaches available in *P. berghei*, including no clear depletion of protein levels upon introduction of an auxin-inducible degradation motif and no recovery of parasites following attempts to integrate blood-stage specific promoters. We therefore interrogated the AKiT1 ortholog in *T. gondii* by fusion with an auxin-inducible degron, speculating that kinetochore architectures with respect to AKiT1 may bear similarities across apicomplexan organisms. Supporting the notion of a conserved role for AKiT1 across the Apicomplexa, tagged protein showed kinetochore-like localizations during *T. gondii* asexual tachyzoite divisions (Fig. 4A), with clearer association of foci with respect to TgNUF2 at kinetochores compared to the centrosome or apicoplast (Fig. S4A).

**Figure 4.**
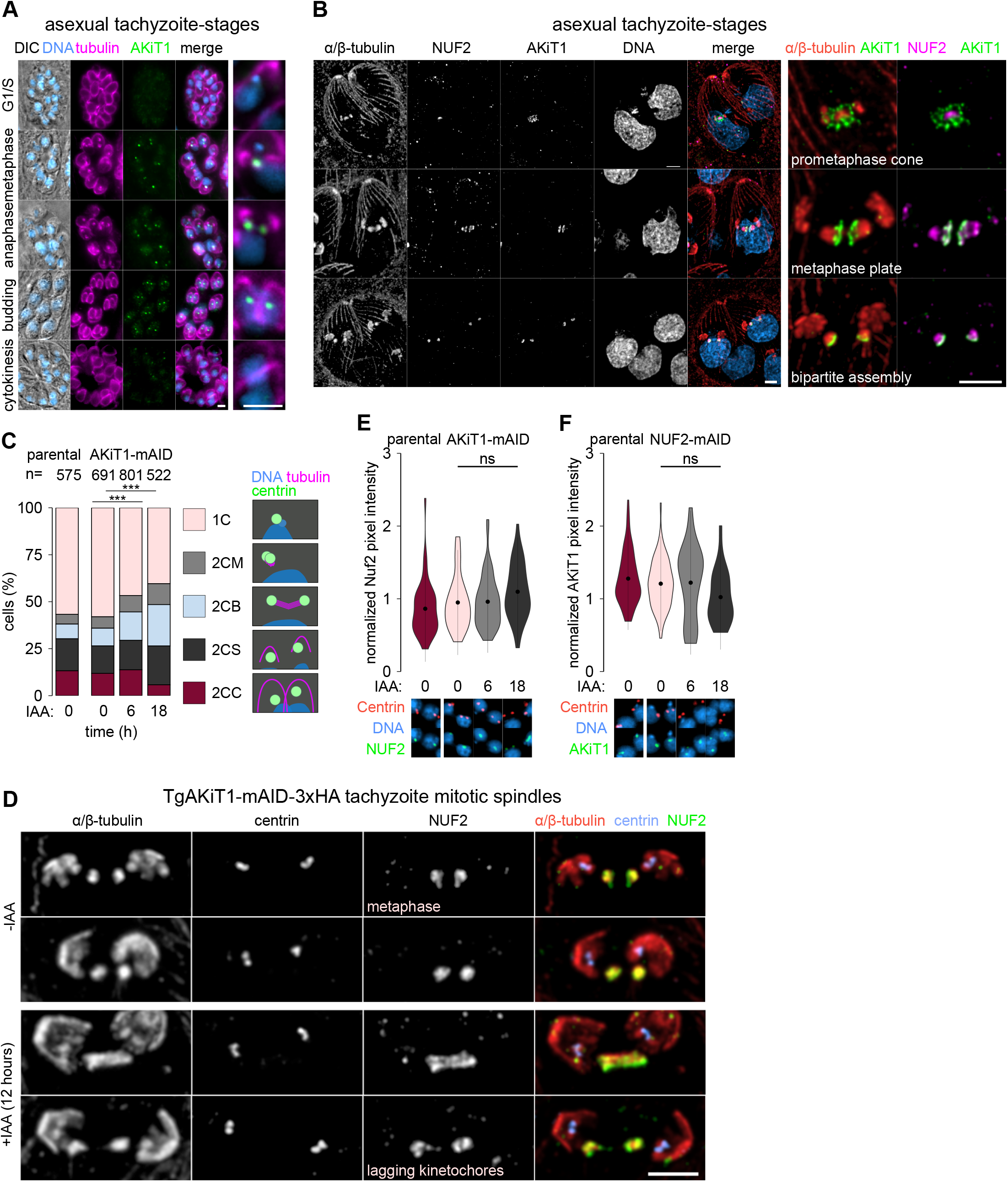
AKiT1 is an essential component of the *Toxoplasma* kinetochore. (A) Micrographs of *T. gondii* tachyzoites expressing AKiT1-2xTy throughout intracellular divisions. Counter-staining of DNA with DAPI (cyan), tubulin (magenta) and DIC images are also shown. Bar: 5 μm. (B) U-ExM revealed AKiT1 at the tachyzoite mitotic spindle (identified by α/β-tubulin counter-stain) and kinetochores (NUF2) throughout mitosis. Counter-staining of DNA with DAPI (cyan) also shown. Bar: 2 μm. (C) Depletion of AKiT1-mAID-3xHA prevented proper mitotic progression due to a buildup of cells with duplicated centrosomes and bipolar spindles (2CB) (**, P < 0.01; ***, P < 0.001; Chi^2^ test). (D) U-ExM further characterized phenotypic changes post-depletion, revealing tachyzoites with stretched mitotic spindles and lagging kinetochores. Bar: 1 µm. (E & F) Levels and localization of NUF2 and AKiT1 kinetochore components in cells at 0, 6 and 18 hours post-depletion of either component. Representative images shown below.

In contrast to the localization of TgSKA2, U-ExM revealed TgAKiT1 foci forming a cone around the tachyzoite prometaphase spindle (Fig. 4B). Little overlap was seen between TgAKiT1 and the majority of TgNUF2 signal at the spindle. This distinction exaggerated upon formation of the diamond spindle with TgAKiT1 forming a striking metaphase pate, comparable to that seen during *Plasmodium* asexual blood-stage division, prior to segregation of foci at anaphase.

As with depletion of TgSKA components, reducing levels of TgAKiT1 (Fig. S4B & C) led to a rapid and severe reduction of intracellular growth (Fig. S4D), producing vacuoles with abnormal accumulations of DNA 18 hours post-depletion (Fig. S4E). TgAKiT1 depletion primarily stalled cells progressing through anaphase, with a greater increase in the number of cells with bipolar spindles (Fig. 4C), resulting in elongated mitotic spindles and aberrant kinetochore segregation (Fig. 4D).

Surprisingly, despite the striking centromere proximal localization of TgAKiT1 relative to TgNUF2, no clear reduction in TgNUF2 levels at the spindle were seen in tachyzoites depleted for TgAKiT1 relative to parental and non-induced controls (Fig. 4E). Similarly, levels of TgAKiT1 at kinetochores was not clearly affected by depletion of TgNUF2 (Fig. 4F), suggesting neither protein is dependent on one another for assembly onto kinetochores.

### An AKiT complex bridges the outer kinetochore to the centromere

In animals and fungi, kinetochores are composed of hierarchical assemblies totaling ∼50 - 100 proteins. Whilst exciting, the addition of AKiT1 increases our repertoire of validated apicomplexan kinetochore components to a mere order of magnitude less than in animal or fungal systems. Streamlining of redundant kinetochore composition is not uncommon in eukaryotes (Przewloka et al., 2007; Westermann & Schleiffer, 2013). Furthermore, the frequency of gene loss compared to gain has been associated with transition to parasitism in the Apicomplexa (Mathur et al., 2019). It’s quite possible few apicomplexan kinetochore proteins manage the roles of many in other organisms. However, neither the centromeric histone variant CENH3 nor SEA1 (a previously reported homolog of CENP-C (Perrin et al., 2021; Verma & Surolia, 2014)) were identified in immunoprecipitates of NUF2 or SKA2, suggesting additional components may bridge the outer kinetochore to the centromere. To test whether PbAKiT1 interacts with centromeric proteins, we employed a similar proximity based approach and affinity purified protein, however with an additional limited cross-linking condition (D’Archivio & Wickstead, 2017). Given that at least in animals the SPC24:SPC25 heterodimer forms a direct interaction with the CCAN, we additionally immunopurified PbSPC24 under the same conditions, to compare interactors (Fig. 5A). Further supporting the identification of a biochemically stable outer kinetochore complex, a clear reciprocal enrichment was seen for each component of NDC80/NUF2 across all immunoprecipitates of SPC24. Additionally, PbAKiT1 was enriched – albeit marginally – upon cross-linking, reinforcing a less stable interaction between SPC24 and PbAKiT1 compared to SPC24 and other NDC80/NUF2 components. Two additional proteins of unknown function (Gene IDs: PBANKA_1243900, PBANKA_0522000) were enriched to comparable levels. Proteins belonging to the pre-replicative complex MCM6, ORC1 and CDC6 were enriched upon higher cross-linking conditions and this set also contained components of the SKA complex and STU2, along with six additional hypothetical proteins of unknown function. SEA1 was identified upon high cross-linking alone.

**Figure 5.**
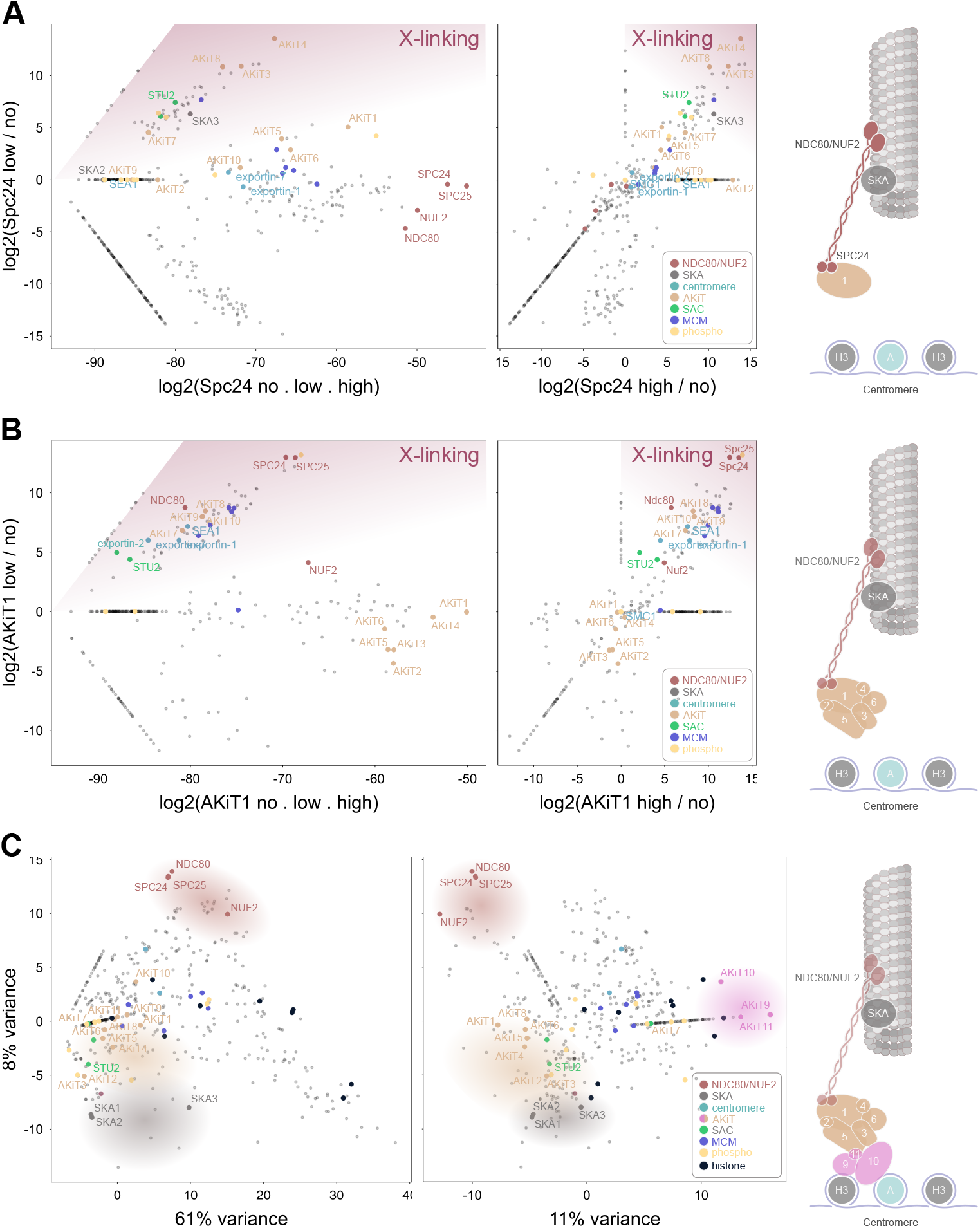
The *Plasmodium* kinetochore is composed of at least four biochemically stable compartments. Relative enrichment of proteins immuno-purified with SPC24-3xHA (A) and AKiT1-3xHA under conditions of “low” compared to “high” and “no” cross-linking conditions (left), or “low” compared to “high” cross-linking (right). (C) Relative enrichment of proteins immuno-purified (without cross-linking) with NUF2, SPC24, AKiT1, AKiT8, AKiT9, SKA2 and PBANKA_1343200, in addition to spindle controls; NEK1 and KIN8X, displayed in principal components 1 and 3 (left) and 2 and 3 (right). Schematic interpretations are shown on the far right. Intensities for all 780 proteins detected across experiments are presented in Table S3.

Along with AKiT1, 5 proteins of unknown function were most abundant across PbAKiT1 immunoprecipitates (Fig. 5B), 4 of which (Gene IDs: PBANKA_0612200, PBANKA_1310500, PBANKA_1243900 and PBANKA_0522000) were also enriched in purifications of SPC24 upon cross- linking. In addition to components of the pre-replicative complex, MCM2 - 7, and STU2, proteins previously reported to be at the apicomplexan centromere SMC1, Exportin 1 and Exportin 7 (Francia et al., 2020) in addition to SEA1 were enriched upon low cross-linking conditions, and this threshold showed reciprocal enrichment for all components of the NDC80/NUF2 complex.

Together, these data suggest the existence of biochemically stable complex that includes PbAKiT1 that deposits between NDC80/NUF2 complex at the spindle and proteins at the centromere. Each tagged- hypothetical protein showed PbAKiT1-like localization patterns (Fig. S5A & C), colocalizing with NUF2 during blood-stage and microgametocyte mitosis, also present as foci in activated female macrogametes and accumulating as four puncta in fully developed ookinetes. We therefore named these proteins AKiT2 - 6.

Interestingly, three proteins of unknown function (Gene IDs: PBANKA_0612300, PBANKA_0213200 and PBANKA_1307000) formed a distinct enrichment profile compared to AKiTs 1 - 6 across immunoprecipitations (Fig. 5A & B). Each tagged protein showed characteristic kinetochore localizations (Fig. S5B & C), with that encoded by PBANKA_0612300 displaying an additional localization reminiscent of the nuclear membrane during sexual development. We named these proteins AKiT7 - 9, respectively. To test the extent and composition of additional complexes at the malaria parasite kinetochore, we similarly affinity purified tagged protein. The relative abundances of co-purifying proteins identified in immunoprecipitations of *P. berghei* kinetochore proteins (without cross-linking) were then assessed by principal component analysis (Fig. 5C) (Brusini et al., 2021), including NUF2, SPC24, SKA2, AKiT1, AKiT7, AKiT8 and AKiT9, alongside controls previously shown to localize in the vicinity of the *Plasmodium* spindle, KIN8X and NEK1 (Dorin-Semblat et al., 2011; Zeeshan et al., 2019). Principal components 1 and 3 encompass 69% of the total variance in the data and show distinct clustering of NDC80/NUF2 and SKA components relative to all AKiTs. Principle components 2 and 3 further resolved AKiT clustering, with the distinction of AKiTs 1 - 6 relative to AKiT9, itself clustering with histones H3 and H4, the histone modifier SPT16, and two additional proteins of unknown function (Gene IDs: PBANKA_0406000 and PBANKA_0803900) that we named AKiT10 and AKiT11, respectively.

Taken together, our immuno-affinity purification strategy has identified 4 biochemically stable compartments at the *Plasmodium* kinetochore (Table S4). Between compartments, more labile interactions were also detected between the outer kinetochore NDC80/NUF2 complex and PbAKiT1, itself closely associated with 5 additional AKiT proteins. AKiT1 - 6 also interact with AKiT9 - 11, which, given their association with proteins known to interact with DNA, likely form the most centromere proximal kinetochore sub-domain identified in this study.

### Plasmodium kinetochores include distant relatives of CENP-C and the SAC

AKiTs 1 - 11 have not previously been annotated with protein function. Furthermore, sequence analysis of AKiT proteins revealed no clearly predicted Pfam domains (Fig. 6A), excepting an unusual allancoitase domain in AKiT8, not previously identified in any kinetochore protein, and an AT-hook within AKiT10. Iterative HMM profile-profile comparisons failed to find any evidence that AKiTs 1 - 6, 8 or 11 are similar to known eukaryotic kinetochore components (Fig. S6A - G). In contrast, we detected strong similarity between AKiT7 and the spindle assembly checkpoint protein MAD1 (Fig. 6B & C & S6H). Similarly, scores for AKiT9 and AKiT10, in addition to the previously described SEA1 (Perrin et al., 2021), suggest these genes are homologous to the conserved CCAN component CENP-C (Fig. 6D & E).

**Figure 6.**
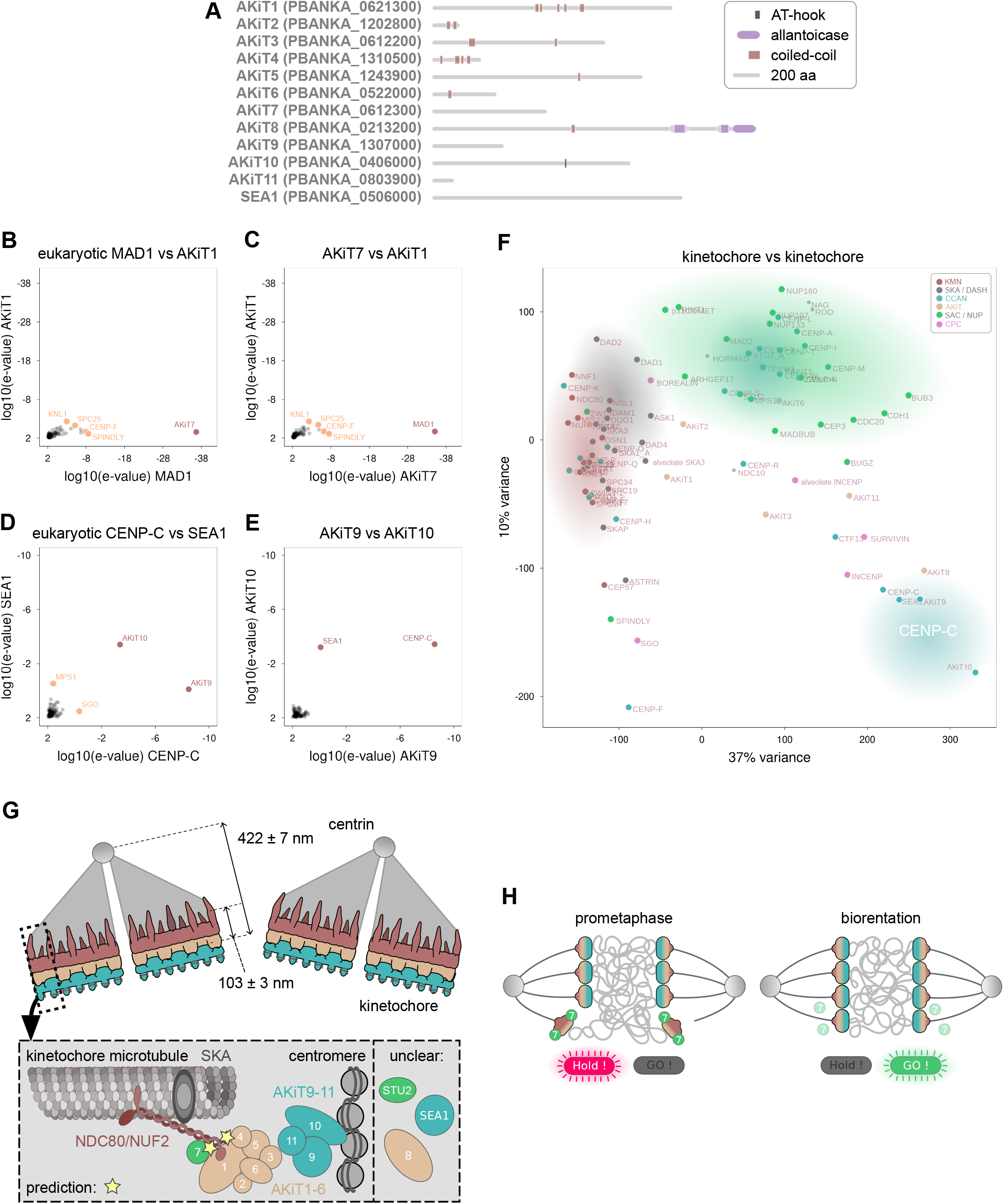
AKiT7, AKiT9 and AKiT10 are related to known eukaryotic kinetochore and spindle assembly checkpoint proteins. (A) Protein architectures, including Pfam domains and coiled-coils, for *P. berghei* homologs of AKiTs 1-11 and SEA1, predicted using HMMer tools (hmmscan). (B-E) Reciprocal HMM profile-profile comparisons using kinetochore HMMs that include alveolate homologs of AKiT7 (PBANKA_0612300), AKiT9 (PBANKA_1307000), AKiT10 (PBANKA_0406000) and SEA1 (PBANKA_0506000) and pan-eukaryotic HMMs identify similarity with MAD1 and CENP-C as highest scoring hits. In red are high confidence/scoring HMMs and in orange low confidence hits. (F) Scores of all kinetochore HMM profile-profile comparisons displayed in principal components of largest variance. Clusters are colored according to previously demonstrated functional (biochemically purified complexes) and/or phylogenetic (sequence similarity) interactions. (G) Schematic model representation of the *Plasmodium* kinetochore. Overall organization based on electron microscopy images, presented previously (Guttery et al., 2012), and molecular architectures combine biochemically purified complexes and localization data presented in this study. (H) Proposed metaphase kinetochore organization with the inclusion of AKiT7/MAD1 as a spindle assembly checkpoint component, based on biochemical and localization data presented in this study.

Similar phylogenetic profiles have previously been shown to reflect and predict functional interactions between eukaryotic kinetochore proteins (van Hooff et al., 2017). To investigate the evolutionary relationships and predict interactions between newly identified AKiT proteins, we assessed profile-profile scores for all kinetochore HMMs by principal components analysis (n = 90, Fig. 6F). In agreement with previously reported homology and functional interaction, we found intracomplex clustering between the SKA (SKA1 - 3), DASH (DUO, DAM, DAD and ASK), CENP-O (CENP-OUPQ), CENP-T (CENP-TWSX), in addition to components of the spindle assembly checkpoint (SAC) with nuclear pore proteins (NUPs) and the KMN network. A cluster of profiles for AKiT9, AKiT10, SEA1 and CENP-C further supports evidence these genes are homologous. AKiT7 clustered within the KMN network, and we predict this protein’s function to be more tightly linked with that of the outer kinetochore and AKiT1 - 6 compartments compared to the CCAN in the malaria parasite (Fig. 6G). We did not find clustering of AKiT proteins, but instead found AKiT4 within KMN and AKiT5 - 6 within SAC clusters, respectively.

Overall, our data indicates that the CCAN in malaria parasites includes at least two distant relatives of CENP-C. Between the CCAN and the outer kinetochore deposits an AKiT1 - 6 compartment, comprised of proteins bearing very little detectable similarity to known kinetochore proteins and patterns of phylogenetic profiles suggest only AKiT4 remotely within the outer kinetochore cluster. AKiT7 is the MAD1 component of the SAC and its function is most likely linked with that of the KMN network.

### AKiT1 requires CENP-C for kinetochore localization in Toxoplasma

Identification of CENP-C within the *Plasmodium* CCAN certainly suggests homologous structures drive assembly of very different kinetochores in these parasites. Interestingly, we identified only a single CENP-C gene in *T. gondii* (Fig. 7A). To investigate whether assembly of the NDC80/NUF2 complex or AKiT1 onto kinetochores relies upon CENP-C in *Toxoplasma*, we similarly tagged protein with the auxin-inducible degron fused to 3xHA, alongside tagged TgAKiT1 and TgNUF2 kinetochore markers. Consistent with a conserved localization to eukaryotic kinetochores, the majority of tagged TgCENP-C localized to the mitotic spindle, closely associated with TgAKiT1 at kinetochores throughout tachyzoite divisions (Fig. 7B & S7B). Furthermore, U-ExM revealed TgCENP-C foci were present on the centromeric side of the metaphase plate relative to TgAKiT1 at the tachyzoite mitotic spindle (Fig. 7C & S7B), maintaining this bipartite localization following anaphase. Supporting the notion that CENP-C forms a platform for kinetochore assembly, recruitment of TgAKiT1 to kinetochore foci was abolished upon depletion of TgCENP-C (Fig. 7D & S7C). Surprisingly however, levels of TgNUF2 at kinetochore foci were unaffected upon depletion of TgCENP-C (Fig. 7E), despite the severe phenotype on parasite proliferation and spindle assembly at both 6 hours and 18 hours post-depletion (Fig. S7D & E).

**Figure 7.**
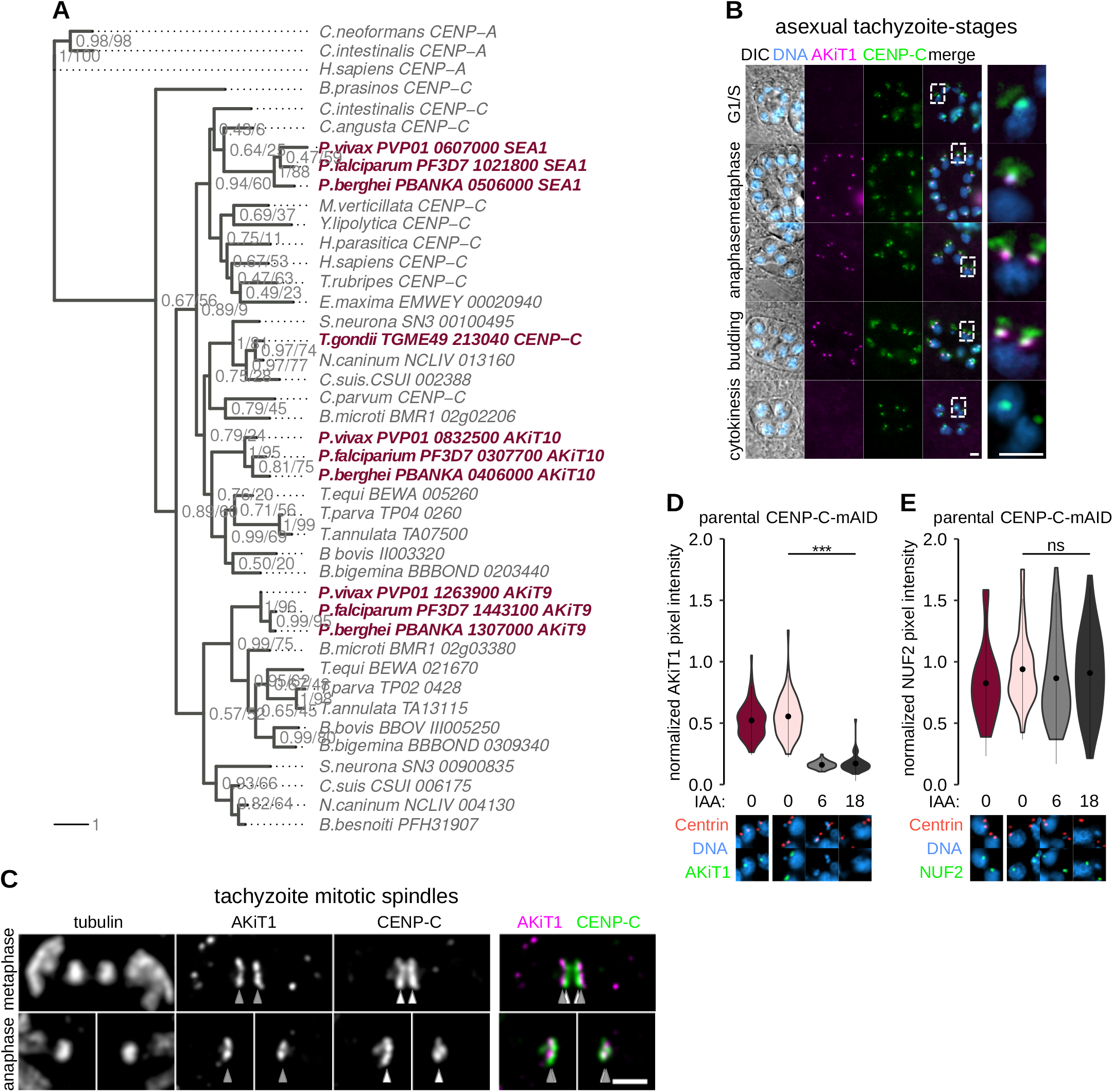
AKiT1 localization to kinetochores is dependent upon CENP-C in *Toxoplasma*. (A) Result of a maximum likelihood inference based on an alignment of sequences retrieved following iterative HMM scans for homologs of *Plasmodium* AKiT9 (PBANKA_1307000), AKiT10 (PBANKA_0406000) and SEA1 (PBANKA_0506000). Numbers beside nodes indicate support from Bayesian posterior probabilities / maximum likelihood-bootstrap values (100 replicates). (B) Micrographs of *Toxoplasma gondii* tachyzoites expressing TgCENP-C fused to mAID-3xHA (green) and AKiT1-2xTy (magenta) throughout intracellular divisions. Counter-staining of DNA with DAPI (cyan) and DIC images are also shown. Bar: 5 μm. (C) U-ExM revealed CENP-C (green) localizes to the centromeric side of the metaphase plate relative to AKiT1 (magenta). Bar: 1 μm. (D & E) Levels and localization of NUF2 and AKiT1 kinetochore components in cells at 0, 6 and 18 h post-depletion of CENP-C, assessed by immunofluorescence against Centrin1 and 2xTy tagged protein. Representative images shown below. DNA stain DAPI.

## Discussion

### Apicomplexan models to study eukaryotic kinetochores

In this study, we have looked into the composition and function of kinetochores in the Apicomplexa, a group of parasites belonging to a lineage that diverged early in eukaryotic evolution from animals and fungi and an obvious niche in kinetochore research. In addition to eukaryotic placement, apicomplexan parasites appear to divide differently to most of the cells of their hosts. A longstanding question remains as to how these parasites maintain fidelity of genome segregation following multiple rounds of DNA replication without concomitant karyokinesis, in spite of an apparent absence of canonical kinetochore and checkpoint proteins. To address these largely unexplored questions we have used two model apicomplexan parasites to gain insights into the conservation of kinetochore proteins and compare protein behavior between lifecycle stages. SKA2 is a highly conserved kinetochore protein that we show localizes to *P. berghei* kinetochores during blood-stage mitosis. However, SKA2 foci were not clearly detected by live fluorescence microscopy during mitosis that occurs at male sexual development and furthermore first appeared at spindle poles before kinetochores during meiosis in the ookinete. Depletion of SKA2 in *T. gondii* asexual tachyzoites led to a strong increase in mitotic index, in particular in cells with bipolar spindles and a metaphase-like arrested state, reminiscent of siRNA-mediated depletion of Ska components in HeLa cells (Hanisch et al., 2006). The significance of differing SKA behavior remains to be explored, however, given the strong requirement for Ska components to enhance spindle attachment in animal cells, it is quite possible kinetochores bind to the spindle with different strengths between apicomplexan lifecycle stages (Helgeson et al., 2018). Low levels of SKA were present at the prometaphase tachyzoite spindle, increasing upon formation of the diamond spindle at metaphase, when kinetochore-microtubule attachments are most likely under greatest tension.

Kinetochore components evolve relatively rapidly compared to most eukaryotic protein sets and become highly specialized along with centromere architectures (Meraldi et al., 2006). Within kinetochores of *Toxoplasma* and *Plasmodium* we have identified a single CENP-C in *Toxoplasma* that is required for assembly of AKiT1 onto kinetochores, whilst *Plasmodium* AKiT9, AKiT10 and SEA1 have been shown to be essential for parasite proliferation in rodent and human malaria parasite models (Table S4) (Bushell et al., 2017; Perrin et al., 2021; Zhang et al., 2018). Notably, centromeres of *P. falciparum* and *P. berghei* are extremely AT-rich compared to *T. gondii* (Brooks et al., 2011; Verma & Surolia, 2018) and AKiT10 encodes an AT-hook found in many DNA-binding proteins (Aravind & Landsman, 1998), including the budding yeast CENP-C homolog, Mif2 (Xiao et al., 2017). Given the diverse modes of division there are likely to be many unique features of the apicomplexan chromosome segregation machinery and f urther comparative studies across the phylum will be needed to confirm whether any AKiT repertoire is truly specific to any organism.

### The molecular architecture of the apicomplexan kinetochore

By combining quantitative proteomics with U-ExM we have identified 4 biochemically stable and distal compartments of the *Plasmodium* kinetochore, including; a three component SKA complex that accumulates at spindle poles and kinetochores; a 6 component AKiT compartment that deposits between the NDC80/NUF2 complex at the outer kinetochore; and the most centromere proximal complex identified formed of AKiT9 - 11. AKiT7, AKiT8 and STU2 also interact with *Plasmodium* kinetochores, however their spatial characterization currently remains elusive. Taken together the findings presented here have provided the first model composition of protein complexes at an apicomplexan kinetochore displayed in Fig. 7G.

How does our model of the apicomplexan kinetochore compare to those in other eukaryotes? All monocentric eukaryotic kinetochores investigated so far are partite, hierarchical assemblies deposited onto specific chromatin environments, themselves often demarked by centromeric nucleosomes (Akiyoshi & Gull, 2014; Cheeseman, 2014; Cortes-Silva et al., 2020; D’Archivio & Wickstead, 2017; Kozgunova et al., 2019). In animals, CENP-C binds directly to CENP-A nucleosomes (Carroll et al., 2010) and interacts with the four-subunit Mis12 complex (Przewloka et al., 2011; Screpanti et al., 2011), itself interacting with KNL1 and the NDC80/NUF2 complex at the spindle (Petrovic et al., 2010). An alternative pathway utilizes the CENP-TWSX complex that bridges DNA to the NDC80/NUF2 complex (Gascoigne et al., 2011). The above design principle is conserved in the Apicomplexa, whose kinetochores we show are also partite hierarchical assemblies. AKiTs 9 - 11 join two identifiable homologs of CENP-C in forming a centromere proximal compartment and likely part of the apicomplexan CCAN. AKiT1 localizes to a midpoint between the CCAN and NDC80/NUF2 complex at the spindle and is dependent upon CENP-C for kinetochore assembly in *T. gondii*, reminiscent of CENP-C binding to Mis12 – KNL-1. However, neither depletion of CENP-C nor AKiT1 in *T. gondii* reduced levels of NUF2 at the spindle and similarly endogenous NUF2 levels were not required for kinetochore recruitment of AKiT1. Recently, a highly elongated *Plasmodium* SPC24 component of the NDC80/NUF2 complex was identified and suggested to bridge the >100 nm distance separating the outer kinetochore from the centromere (Zeeshan et al., 2020). However, this “long form” is poorly conserved across the Apicomplexa. Whilst present in coccidians, it is lacking in *Theileria* and *Babesia*, meaning any direct centromere binding potential would have converged or else been lost by most-closely related hematozoan SPC24 proteins. How the NDC80/NUF2 complex is maintained at the spindle in apicomplexan parasites currently remains unknown, however it is quite possible a pathway similar to CENP-T kinetochore assembly exists or that re-purposing of NDC80/NUF2 function has facilitated binding to microtubules independently of kinetochores.

In human cells, 16 CCAN subunits, forming four sub-complexes are critical for kinetochore assembly and function (Cheeseman, 2014; Guse et al., 2011; Weir et al., 2016). In *Plasmodium*, only three CCAN components, AKiT9 - 11, have been identified. Importantly however, the conserved centromeric histone variant CENP-A that associates with centromeres in *P. falciparum* (Hoeijmakers et al., 2012; Miao et al., 2006; Trelle et al., 2009; Volz et al., 2010) was not found in immunoprecipitates, suggesting the presence of additional as yet unidentified CCAN components in the malaria parasites.

### Origins of the apicomplexan kinetochore

By HMM profile-profile comparisons, we have re-united 6 apicomplexan components identified in this study to a common set, including STU2, SKA2-interacting proteins SKA1 and SKA3, AKiT7, AKiT9 and AKiT10. We did not detect significant similarity between any of AKiTs 1 - 6, which deposit between the CCAN and outer kinetochore, with known eukaryotic kinetochore components. So why is it difficult to identify homology at the apicomplexan kinetochore? Unconventional kinetochores bearing no detectable similarity and arising from a unique origin have been reported in eukaryotic parasites (Akiyoshi & Gull, 2014). Similarly, it is possible AKiTs 1 - 6 are not directly descended from the kinetochores of the last alveolate common ancestor, and in light of the secondary endosymbiosis that forms an integral part of their origins, this scenario is more likely for the Apicomplexa compared to the overall relatively low frequency of horizontal gene transfer in eukaryotes. However, homologs identified in at least some components favors the notion of a common origin for the apicomplexan kinetochore with those in animals and fungi. Gene duplications can be traced in the presence of two INCENP genes (Berry et al., 2018), one of which has been shown to be required for proper mitosis, and we show AKiT9 and AKiT10 are homologous to the conserved kinetochore protein CENP-C. Cupin domains at the C-termini of CENP-C are known to homodimerize (Cohen et al., 2008), and it is tempting to speculate a similar interaction was maintained following duplication that led to formation of a heterodimer. AKiTs 1 - 6 may be more susceptible to change compared to other apicomplexan kinetochore components and this may go some way to explain the lack of similarity detected between this set and all other known eukaryotic kinetochore proteins. Furthermore, identification of AKiT7 as a homolog of the MAD1 component of the spindle assembly checkpoint, further suggests divergence of a KNL1 complex and landing pad for the SAC that we and others have so far been unable to identify (D’Archivio & Wickstead, 2017; Kops et al., 2020; van Hooff et al., 2017). We anticipate that with the ongoing development of greater sensitivity in searches, divergent homologs will be exposed within most kinetochore sets (D’Archivio & Wickstead, 2017; Tromer et al., 2021; Zeeshan et al., 2020).

### Diverse requirements and regulation across apicomplexan modes of division

Fidelity of metazoan chromosome segregation requires the SAC, a surveillance system mediated by the MCC that inhibits CDC20 and activation of the APC/C until proper chromosome bi-orientation. Plasmids containing centromeric sequences are stably maintained throughout the *Plasmodium* lifecycle (Iwanaga et al., 2010, 2012), however the apparent absence of most MCC proteins and the inability to delay cell cycle progression in response to microtubule destabilizing agents has suggested apicomplexan parasites are incapable of a generating a comparable SAC response (Kops et al., 2020; Morrissette & Sibley, 2002). In this study, we reveal alignment of kinetochores during mitosis that occurs during asexual divisions in both *Plasmodium* and *Toxoplasma*, and additionally during meiosis in the ookinete form of the malaria parasite. These architectures are reminiscent of chromosome bi-orientation at the metaphase plate and suggestive of a “hold signal” that prevents precocious entry into anaphase (displayed in Fig. 7H). Furthermore, identification of STU2 at the *Plasmodium* kinetochore hints towards an intrinsic tension-sensing and error-correction mechanism required for establishing bioriented attachments (Miller et al., 2016, 2019). So far, we have been unable to identify a similar metaphase architecture during microgametogenesis. However, kinetochore architecture with respect to tagged-SKA protein localization differs between asexual and sexual stages of division and we speculate this may correlate with requirements to satisfy specific metaphase checkpoints.

Identification of AKiT7 as a homolog of MAD1 and enrichment of protein at kinetochores during nuclear divisions suggest at least a partial SAC response in *Plasmodium* species, and depletion of CDC20 and APC/C components has been shown to prevent proper chromosome condensation during microgametogenesis (Guttery et al., 2012; Wall et al., 2018). Suggestion of any SAC response in the Apicomplexa raises some important questions regarding the existence of a MCC, its recruitment to kinetochores and whether it enables a checkpoint comparable to that in other organisms. Kinases resembling Monopolar Spindle 1 and Polo-Like that are required for MCC assembly onto unattached kinetochores in animal and fungal models have not been detected in the Apicomplexa. Instead, a number of kinases distantly related to Aurora and Cyclin-dependent kinase families have been shown to be implicated in cell cycle control, in particular DNA synthesis and mitosis in both *Plasmodium* and *Toxoplasma* (Balestra et al., 2020; Morahan et al., 2020). Whether phosphoregulation has been repurposed at apicomplexan kinetochores and the effects such divergences have on chromosome segregation will likely broaden our views on the malleability of eukaryotic mitotic checkpoint control.

## Supplementary figures

**Figure S1.**
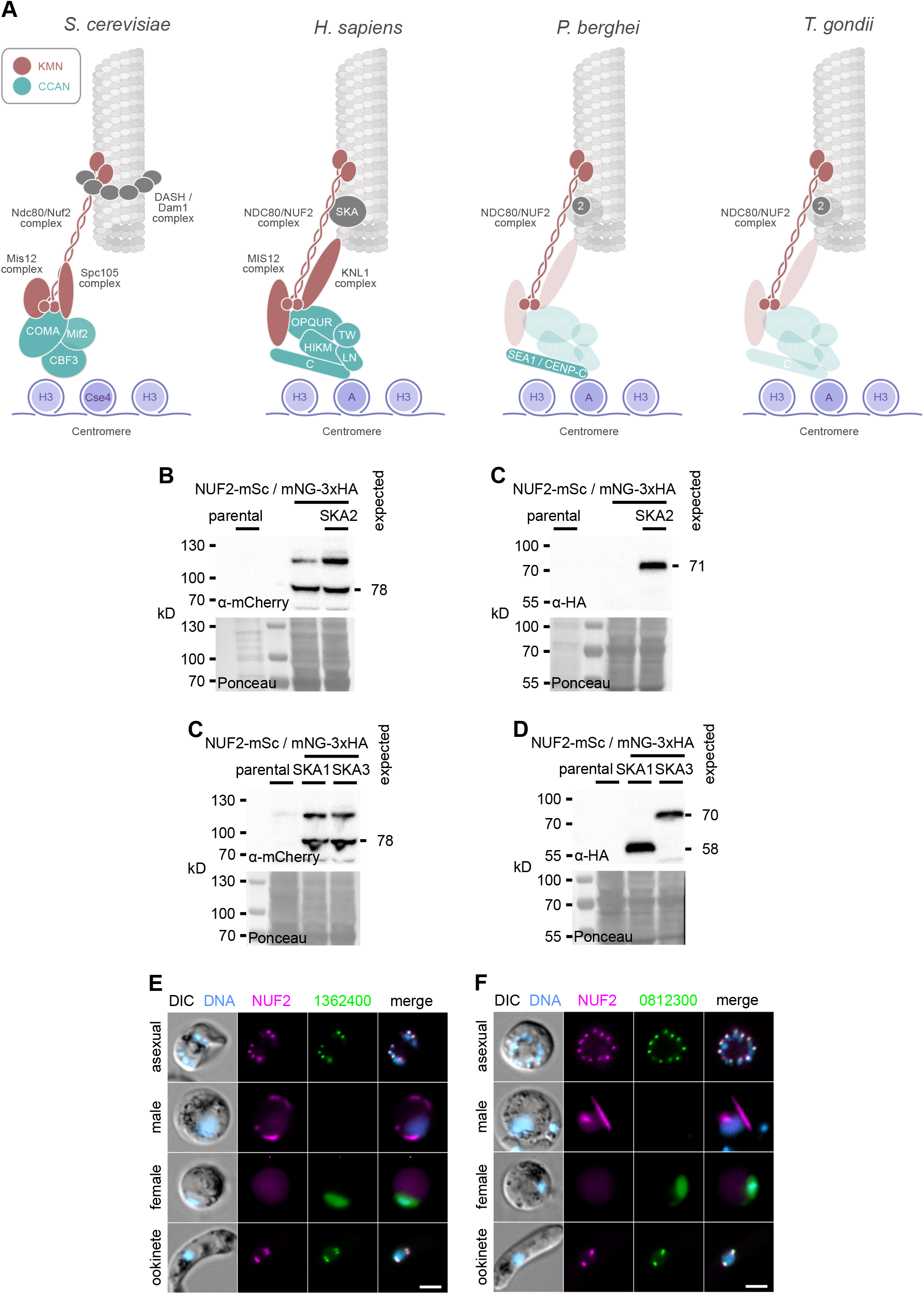
Generation and validation of tagged NUF2 and SKA components in *Plasmodium*. (A) Presence/absence (bold/grey) of kinetochore proteins from model organisms in *Plasmodium* and *Toxoplasma.* Whilst most known components are apparently missing, the NDC80/NUF2 complex and SKA2 are present. (B-D) Immunoblots of malaria parasites expressing tagged kinetochore proteins, probed with either polyclonal α-mCherry or monoclonal α-HA antibodies. Protein loading is shown by Ponceau S stain. (B) Live native fluorescence validated SKA2-interacting proteins tagged with mNG- 3xHA localize to kinetochore foci during asexual blood stages and sexual mosquito stages of development. Counter-staining of DNA with Hoechst 33342 (cyan) and DIC images are also shown. Bar: 2 μm.

**Figure S2.**
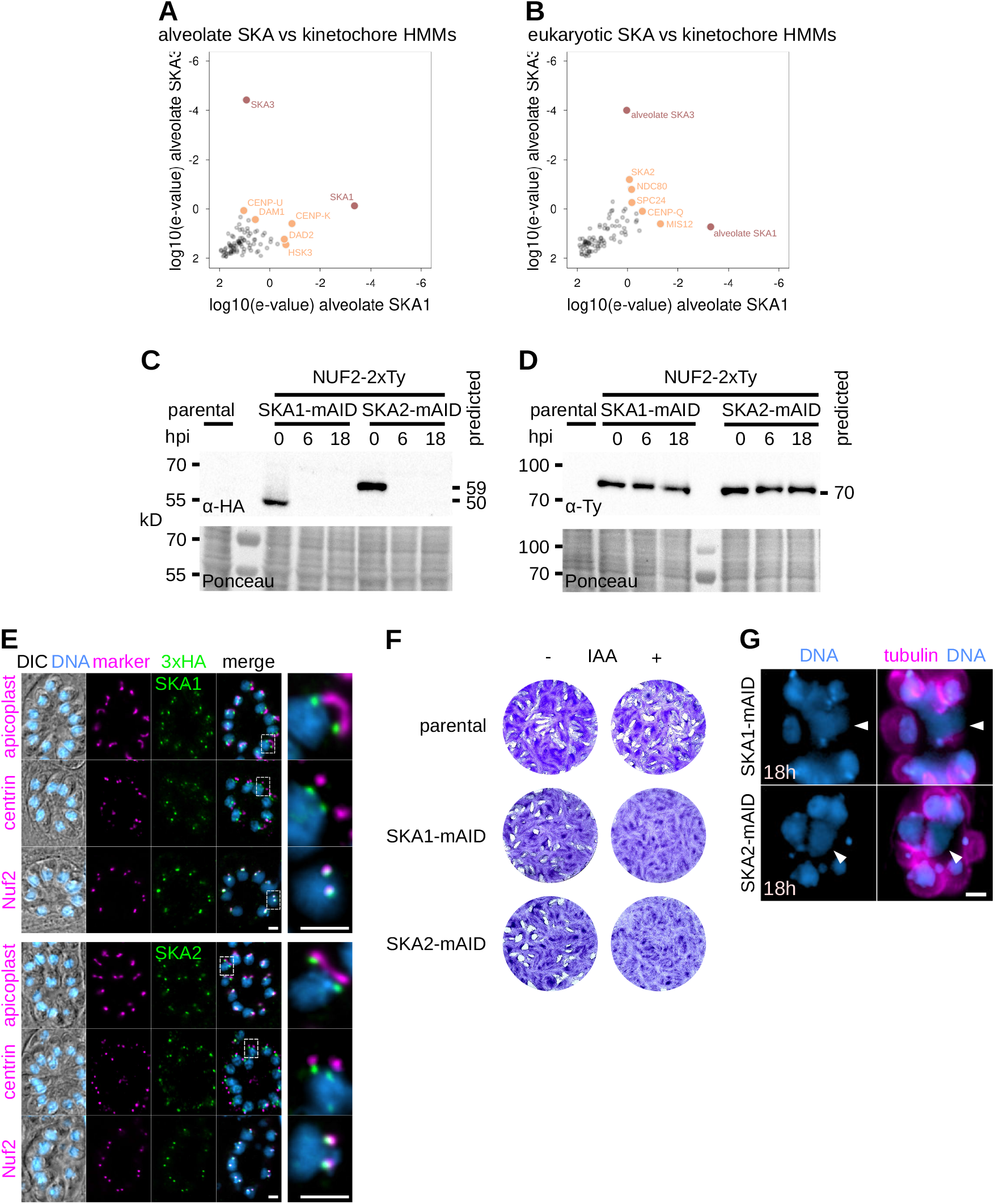
Generation and validation of tagged NUF2 and SKA components in *Toxoplasma*. (A & B) Reciprocal HMM profile-profile comparisons using kinetochore HMMs, including alveolate homologs of SKA2-interacting proteins PBANKA_0812300 and PBANKA_1362400 and pan-eukaryotic SKA1 and SKA3 HMMs, identify one another as highest scoring hits, respectively. In red are high confidence/scoring HMMs and in orange low confidence hits. (C & D) Immunoblots of *T. gondii* parasites expressing tagged kinetochore proteins and showing depletion of mAID-3xHA tagged protein upon induction of auxin. Protein loading is shown by Ponceau S stain. (E) Micrographs of fixed immunofluorescence in *T. gondii* tachyzoites expressing tagged SKA1 and SKA2 throughout intracellular divisions. Counter-staining with antibodies raised against organelle markers (magenta) for the apicoplast (CPN60), centrosome (Centrin1) and kinetochores (NUF2-2xTy). DNA staining with DAPI (cyan) and DIC images are also shown. Bar: 5 μm. (F) Tachyzoites depleted for SKA1 or SKA2 tagged with a mini auxin-inducible degron (mAID-3xHA) failed to form lysis plaques 7 days post-inoculation compared to parental controls. (G) Intracellular vacuoles containing accumulations of DNA (arrow) and no associated cell body were present 18 hours post-depletion of SKA1 or SKA2 tagged with mAID-3xHA. Bar: 5 µm.

**Figure S3.**
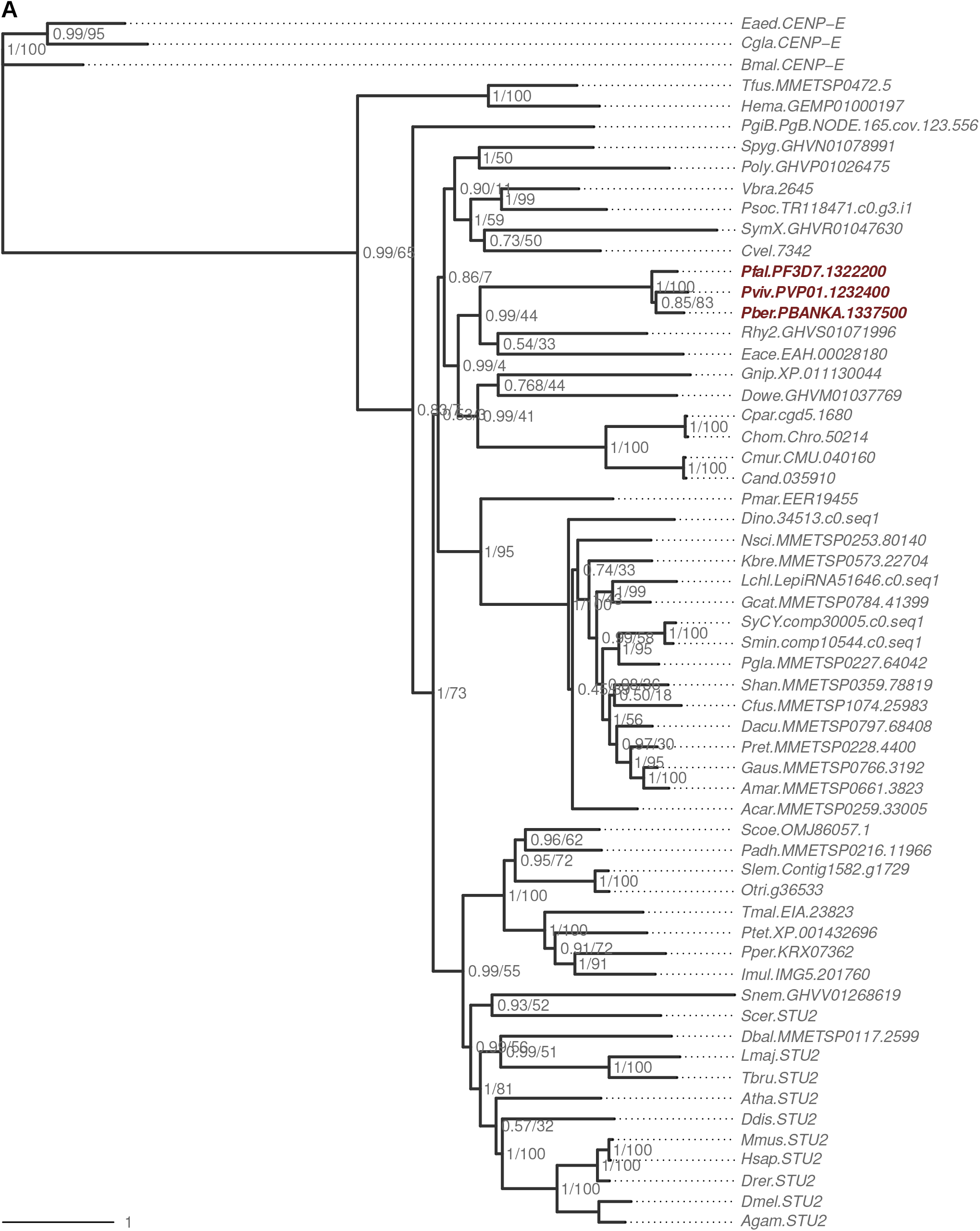
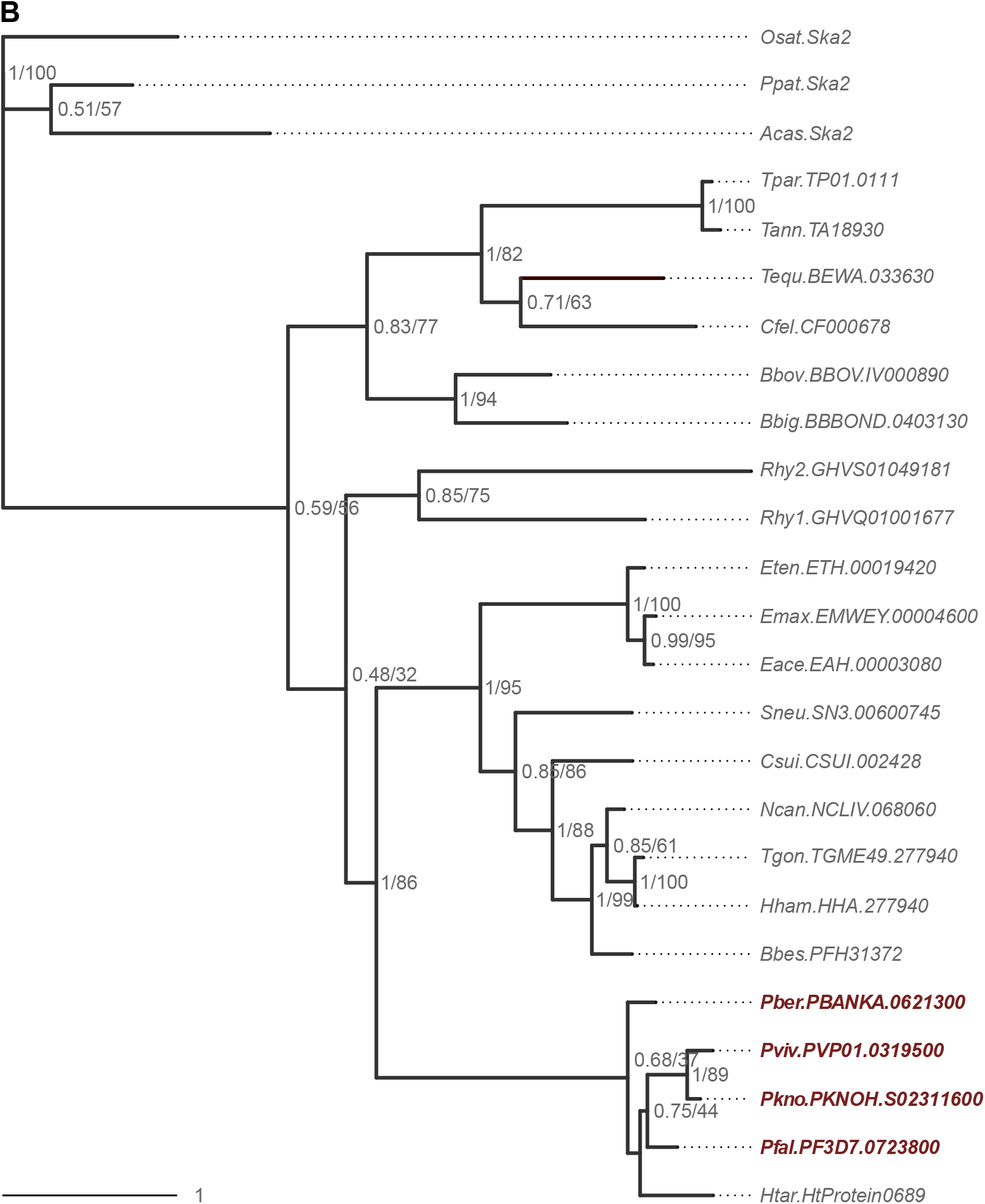
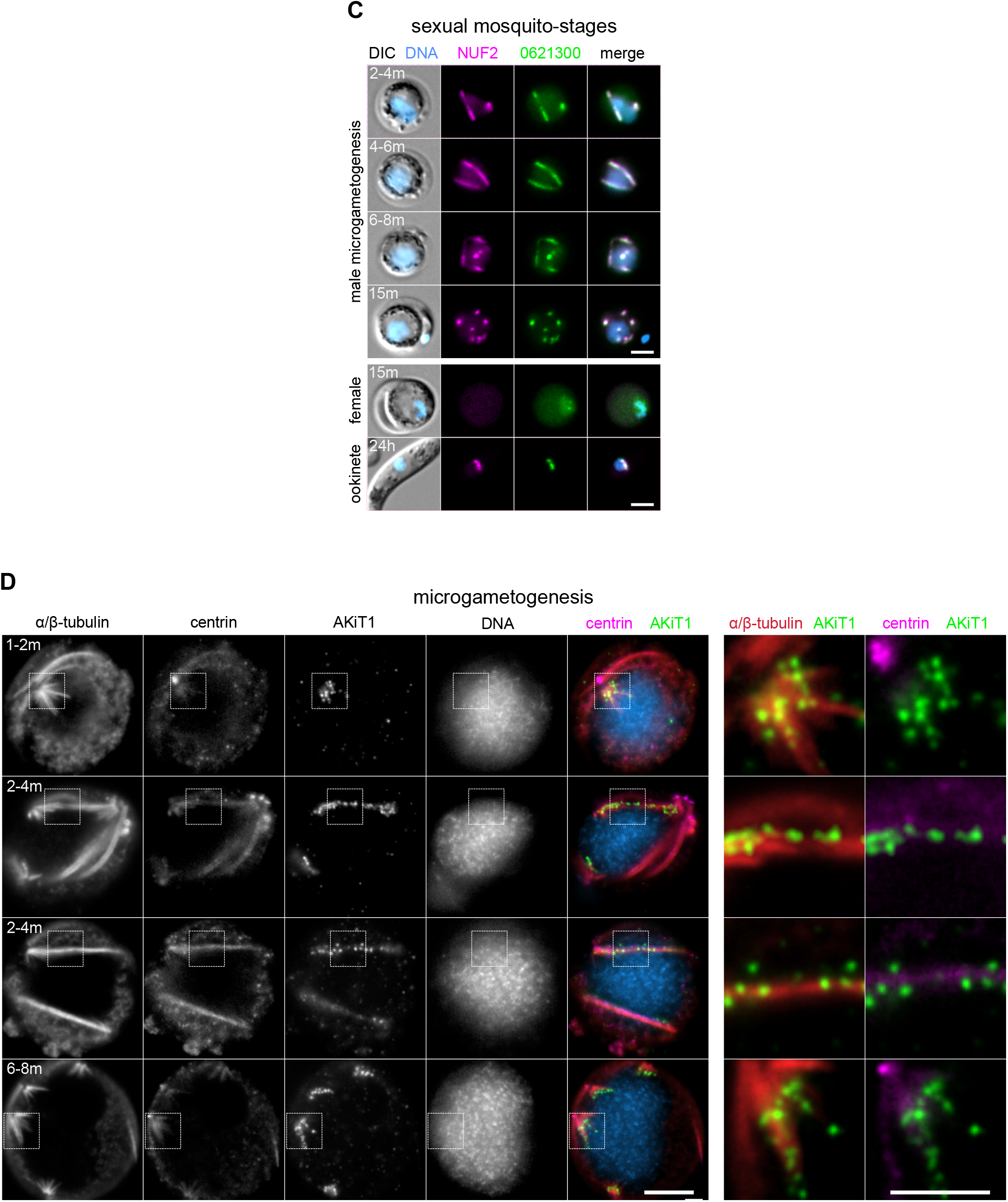
Cross-linking IP of NUF2 identifies novel *Plasmodium* kinetochore proteins. Result of a maximum likelihood inference based on an alignment of sequences retrieved following iterative HMM scans for homologs of STU2 (PBANKA_1337500; A) and AKiT1 (PBANKA_0621300; B). Numbers beside nodes indicate support from Bayesian posterior probabilities / maximum likelihood-bootstrap values (100 replicates). (C) Micrographs of live native fluorescence in malaria parasites expressing tagged kinetochore components NUF2-mScarlet-I (magenta) and AKiT1-mNeonGreen-3xHA (green) during sexual mosquito-stages of development. Counter-staining of DNA with Hoechst 33342 (cyan) and differential interference contrast images (DIC) are also shown. Bar: 2 μm. (D) Widefield micrographs of malaria parasites expressing tagged kinetochore components NUF2-mScarlet-I (magenta) and AKiT1- mNeonGreen-3xHA (green) during microgametogenesis. U-ExM revealed AKiT1 along the spindle (identified by α/β-tubulin counter-stain) and at spindle poles (centrin). Counter-staining of DNA with DAPI (cyan) also shown. Bar: 2 μm.

**Figure S4.**
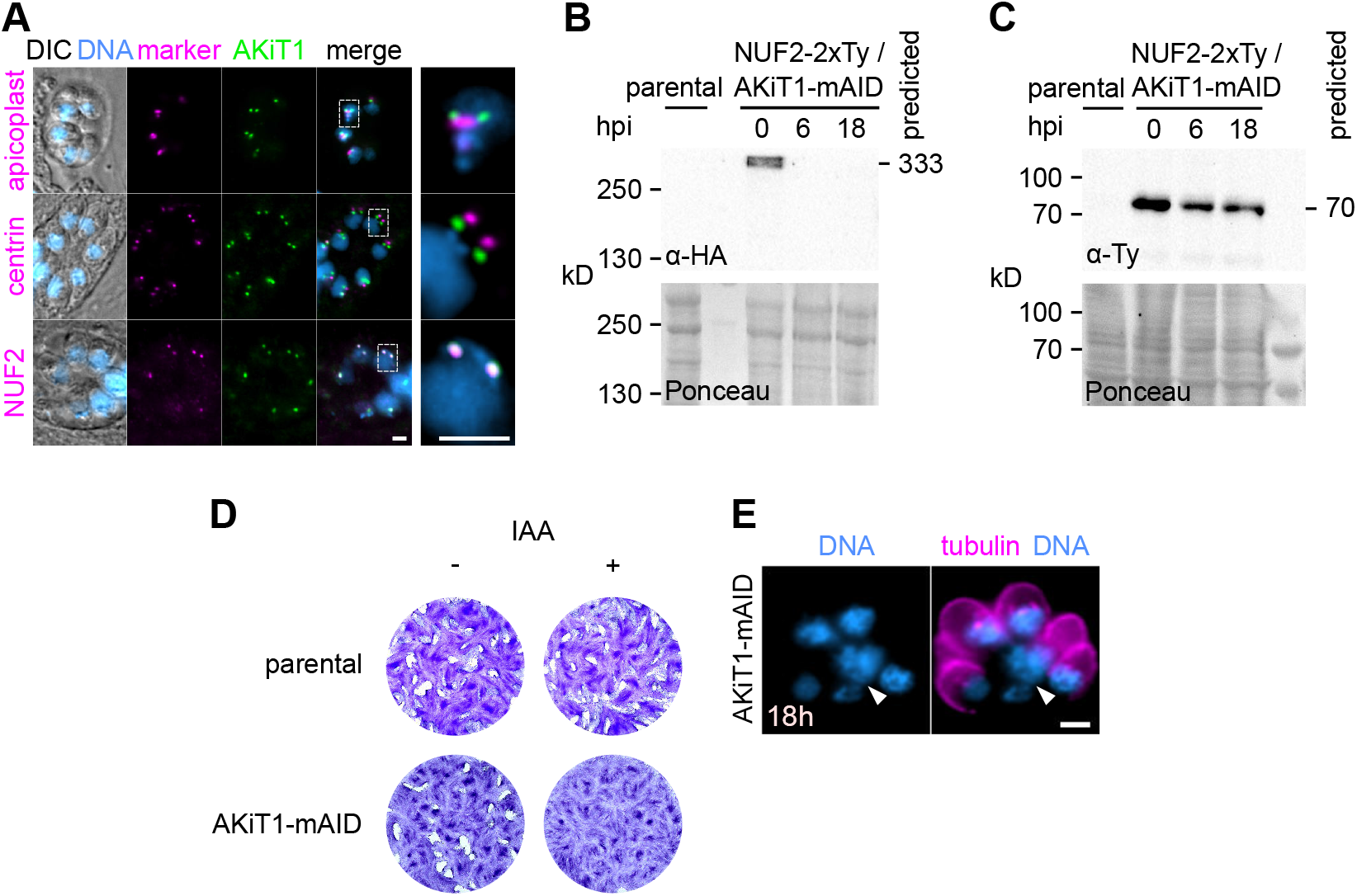
AKiT1 is required for kinetochore segregation in *Toxoplasma gondii*. (A) Micrographs of fixed immunofluorescence in *T. gondii* tachyzoites expressing AKiT1-mAID-3xHA throughout intracellular divisions. Counter-staining with antibodies raised against organelle markers (magenta) for the apicoplast (CPN60), centrosome (Centrin1) and kinetochores (NUF2-2xTy). DNA staining with DAPI (cyan) and DIC images are also shown. Bar: 5 μm. (B & C) Immunoblots of *T. gondii* parasites expressing tagged kinetochore proteins and showing depletion of tagged AKiT1 protein upon induction of auxin. Protein loading is shown by Ponceau S stain. (D) Depletion of AKiT1-mAID-3xHA prevented proper formation of lysis plaques 7 days post-inoculation compared to parental controls. (E) Intracellular vacuoles containing abnormal accumulations of DNA (arrow) were present 18 hours post-depletion of AKiT1-mAID-3xHA. Bar: 5 µm.

**Figure S5.**
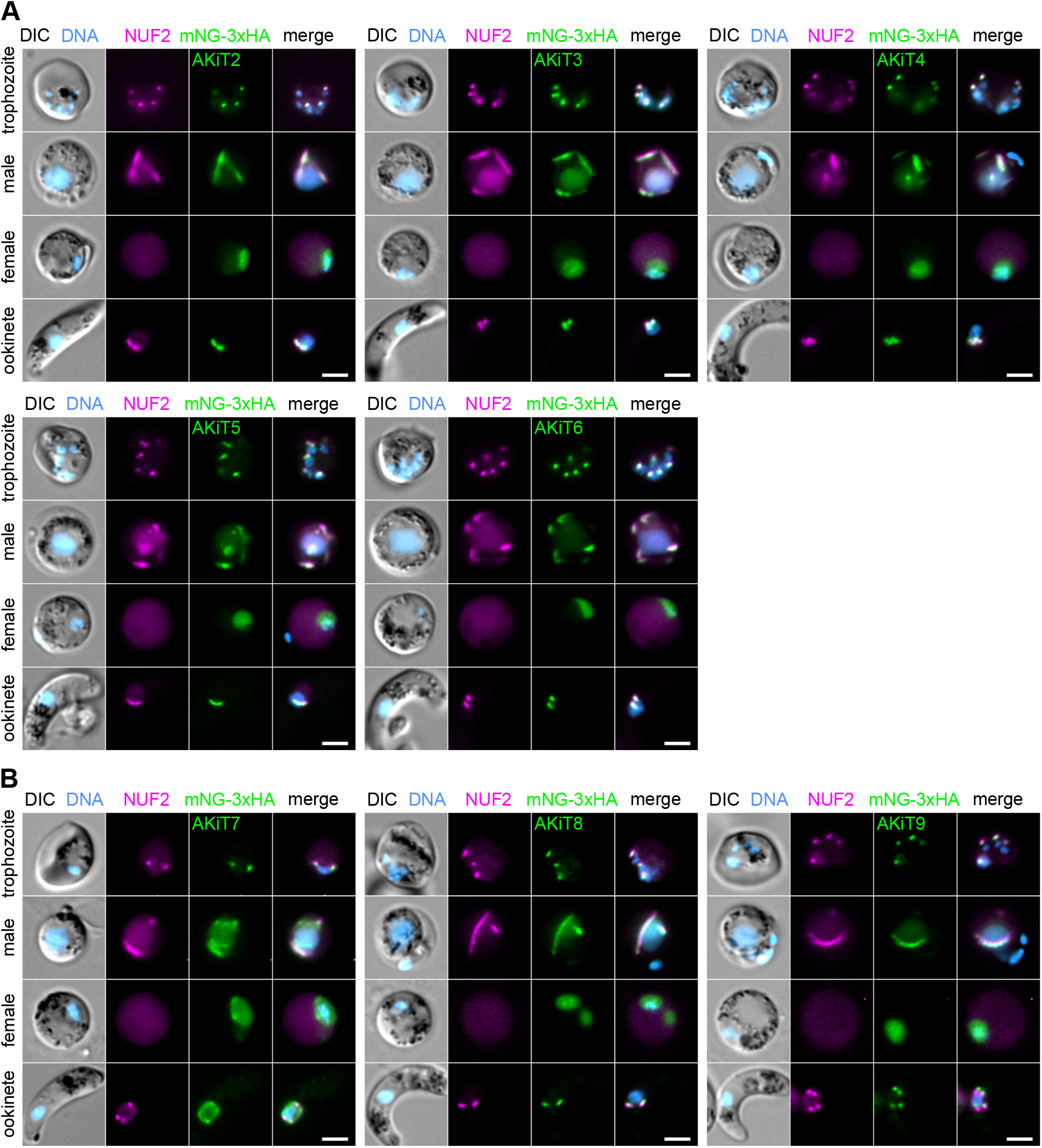
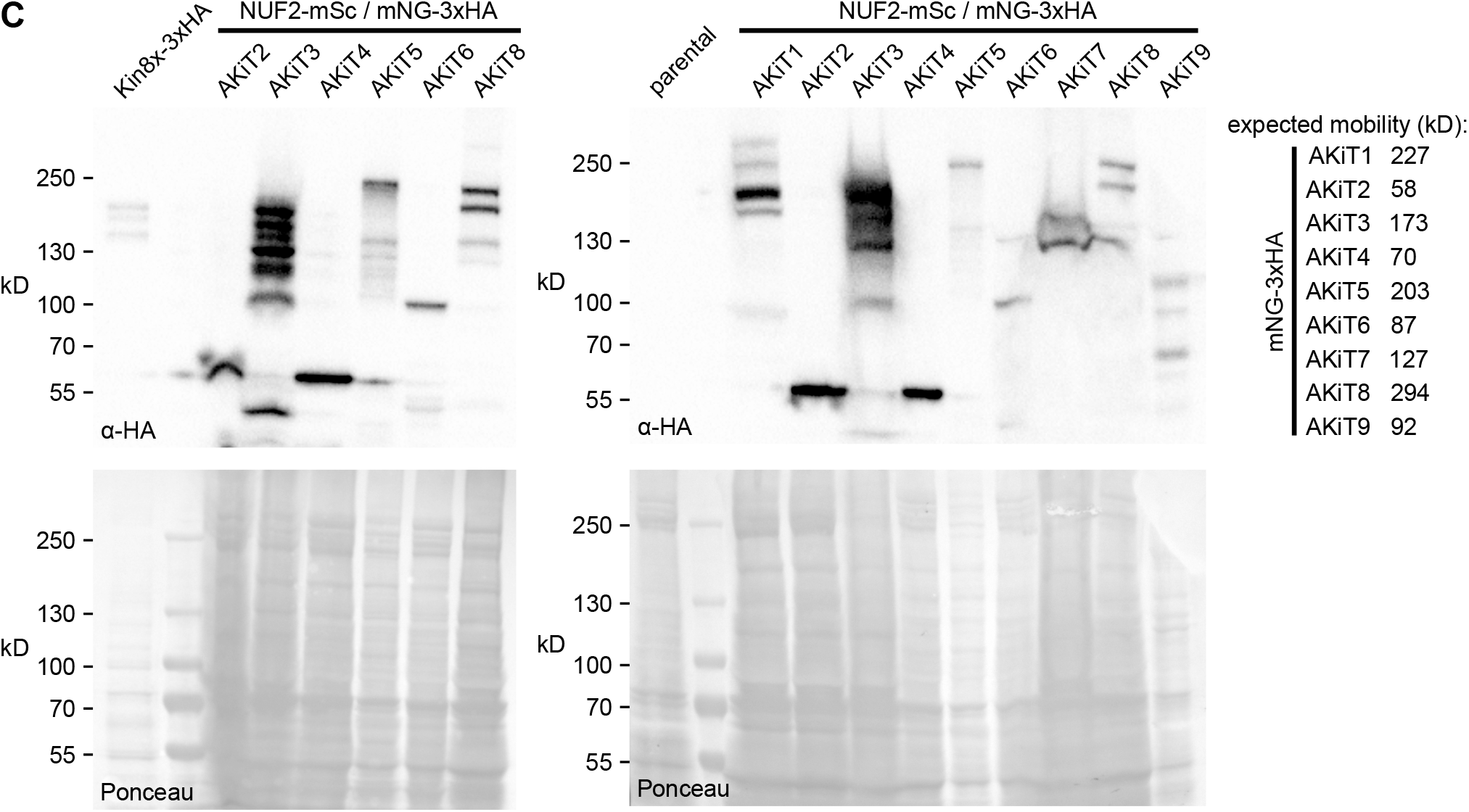
AKiTs are novel identified components of the *Plasmodium* kinetochore. Micrographs of live native fluorescence in malaria parasites expressing NUF2-mScarlet-I (magenta) and tagged AKiTs 1 - 6 (A) and AKiTs 7 - 9 (B) with mNG-3xHA (green) at proliferative stages of the lifecycle. Counter-staining of DNA with Hoechst 33342 (cyan) and differential interference contrast images (DIC) are also shown. Bar: 2 μm. (B) Immunoblots of malaria parasites expressing tagged AKiT proteins, probed with a monoclonal α-HA antibody. Protein loading is shown by Ponceau S stain.

**Figure S6.**
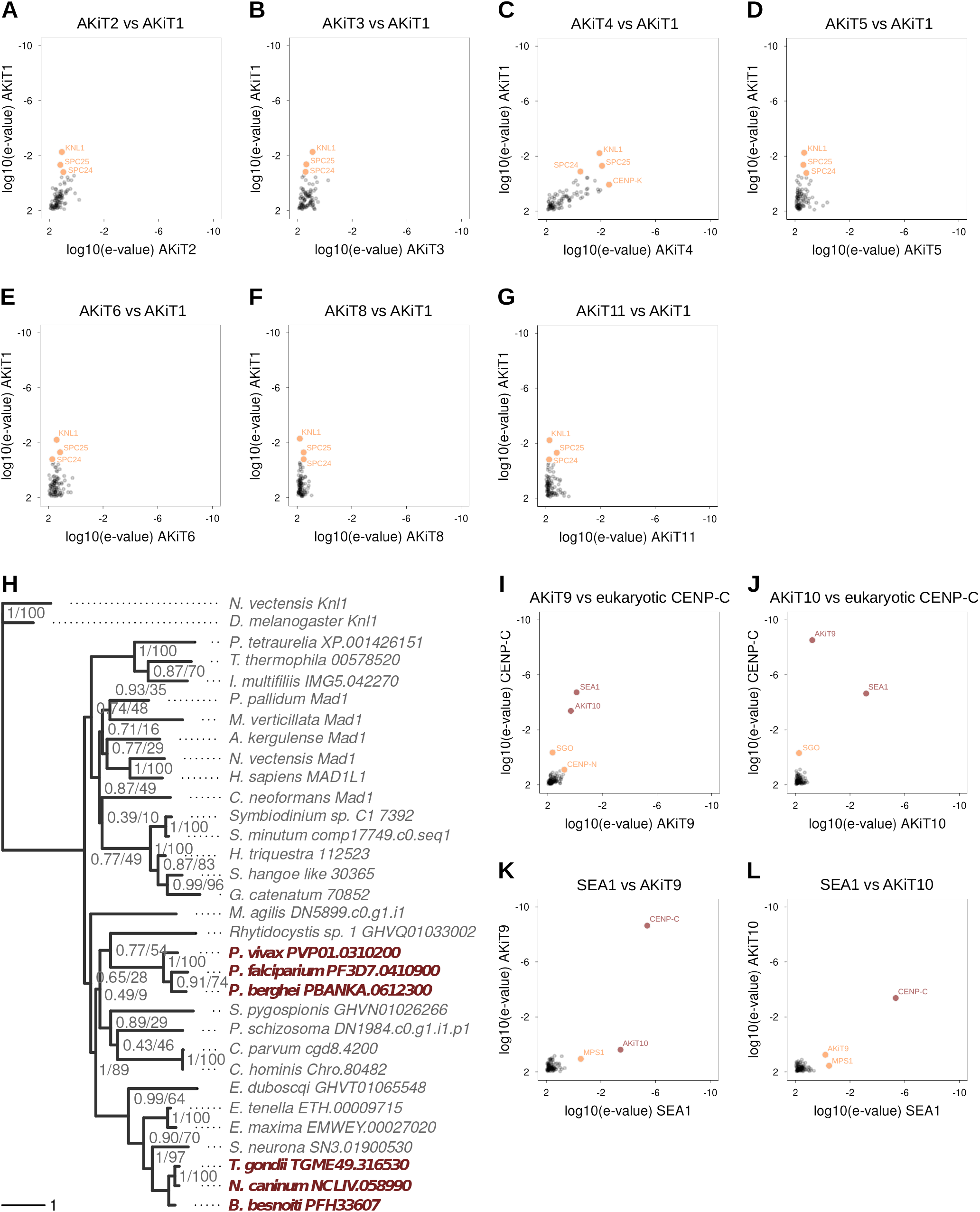
Most AKiTs are not similar to known kinetochore components. (A - G & I - L) HMM profile-profile comparisons using kinetochore HMMs that include alveolate homologs of AKiT proteins and using against AKiT1 as a control set. In red are high confidence/scoring HMMs and in orange low confidence hits. (H) Result of a maximum likelihood inference based on an alignment of sequences retrieved following iterative HMM scans for homologs of *Plasmodium* AKiT7 (PBANKA_0612300). Numbers beside nodes indicate support from Bayesian posterior probabilities / maximum likelihood- bootstrap values (100 replicates).

**Figure S7.**
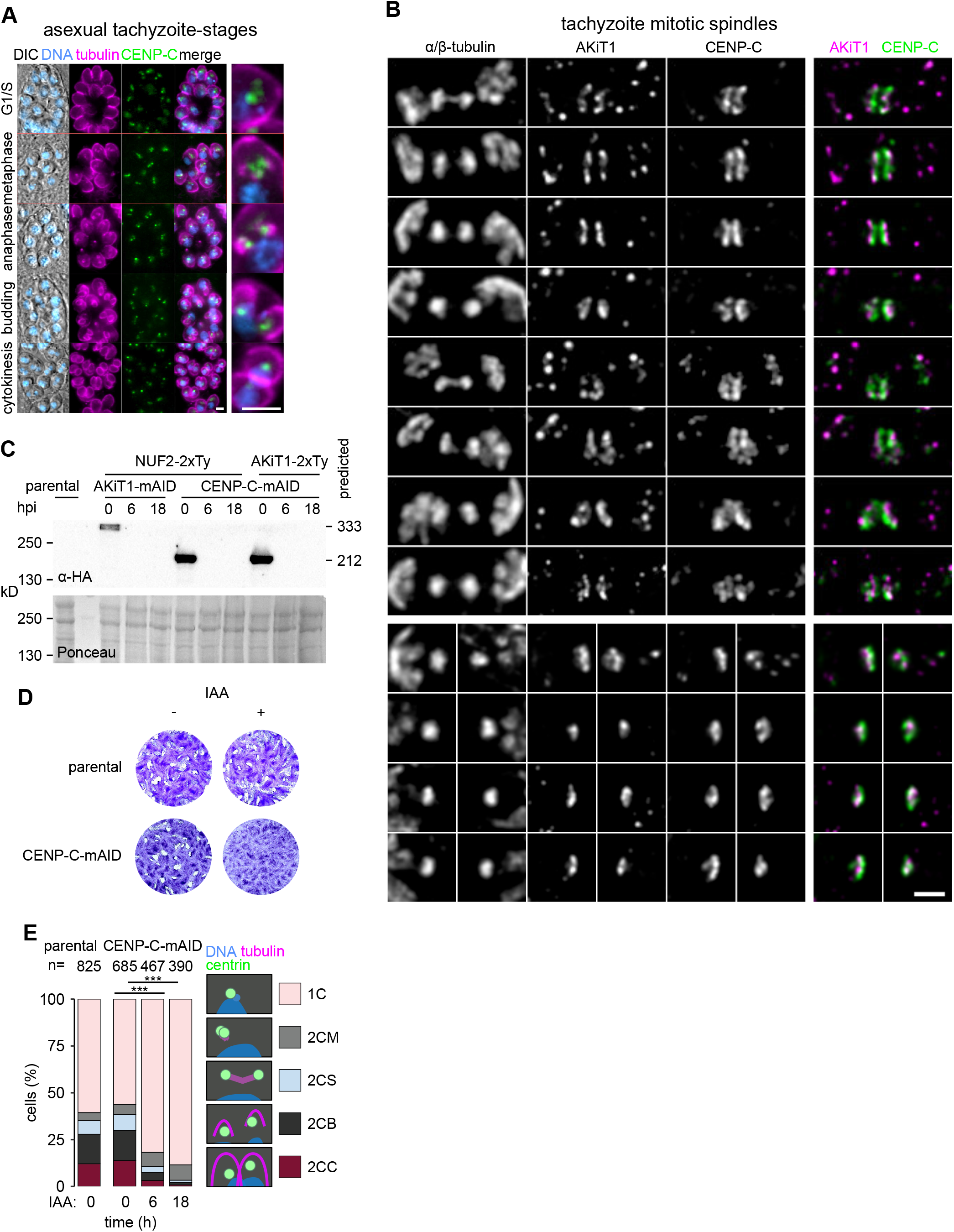
CENP-C is required for *T. gondii* parasite proliferation. (A) Micrographs of fixed immunofluorescence in *T. gondii* tachyzoites expressing tagged CENP-C-mAID-3xHA throughout intracellular divisions. Counter-staining of tubulin (magenta) and DNA staining with DAPI (cyan), in addition to DIC images are also shown. Bar: 5 μm. (B) U-ExM revealed tagged CENP-C (green) localizes to the centromeric side of the metaphase plate relative to AKiT1 (magenta). Bar: 1 μm. (C) Immunoblots of *T. gondii* parasites expressing tagged kinetochore proteins and showing depletion of tagged AKiT1 and CENP-C protein upon induction of auxin. Protein loading is shown by Ponceau S stain. (D) Tachyzoites depleted for CENP-C-mAID-3xHA failed to form lysis plaques 7 days post-inoculation compared to parental controls. (E) Morphological analysis of cells depleted for CENP-C- mAID-3xHA shows a buildup of cells arrested at the onset of mitosis (1C), compared to cells with duplicated centrosomes and monopolar spindles (2CM), bipolar spindles (2CB), segregated centrosomes into daughter cell caps (2CS) and cells in cytokinesis (2CC), assessed by immunofluorescence against Centrin1 (green) and tubulin (magenta) (**, P < 0.01; ***, P < 0.001; Chi ^2^ test).

**Table S1. Sources for detection of putative homologs of apicomplexan kinetochore proteins.** Strategy and nomenclature follows as described (Koreny et al., 2021).

**Table S2. Oligonucleotide primers used for generation of *P. berghei* and *T. gondii* transgenic parasites.**

**Table S3. Label-free semi quantitative mass spectrometry of *P. berghei* kinetochore proteins.**

**Table S4. Apicomplexan kinetochore proteins described in this study.** Essentiality determined by mutagenesis screens deposited on PlasmoDB, for rodent and human malaria parasites (Bushell et al., 2017; Zhang et al., 2018), and ToxoDB, for *T. gondii* (Sidik et al., 2016). *Toxoplasma* protein subcellular compartmentalizations determined by HyperLOPIT (Barylyuk et al., 2020) are also shown.

**Table S5. Antibodies for immunodetection and subcellular localization studies of tagged AKiTs.**

## Author Contributions

Conceptualization: LB, MB. Methodology: LB. Software: LB. Formal analysis: LB. Investigation: LB, NDSP. Original draft preparation: LB. Review and editing: LB, NDSP, MB, DSF. Supervision: MB. Funding acquisition: MB, DSF. All authors contributed to the article and approved the submitted version.

## Acknowledgments

Mass spectrometry and initial quantitation were performed at the Proteomics Core Facility, University of Geneva (www.unige.ch/medecine/proteomique). We thank in particular the service of Alexandre Hainard, Patrizia Arboit and Carla Pasquarello Mosimann in preparation and running of mass spectrometry samples. Microscopy was performed at the Bioimaging core facility, University of Geneva (www.unige.ch/ medecine/bioimaging/en/bioimaging-core-facility), and we thank for the technical assistance of Olivier Brun and François Prodon. We thank Natacha Klages for technical assistance. We also thank Bill Wickstead and Patrick Meraldi for general discussions and critical reading of the manuscript. This work was supported by the Swiss National Science Foundation grant 31003A_179321 MB and 310245753030_185325 DSF. MB is an INSERM and EMBO young investigator.

## Data and materials availability

All data needed to evaluate the conclusions in the paper are present in the paper and/or the supplementary materials. Additional data or reagents are available from authors upon request. Correspondence and requests for materials should be addressed to LB and MB.

## Methods

### Ethics statement

All animal experiments were conducted with the authorization numbers AB_GE18, according to the guidelines and regulations issued by the Swiss Federal Veterinary Office.

### Generation of transgenic parasites targeting constructs

The oligonucleotides used to generate transgenic parasite lines are in Table S2.

For C-terminal tagging of *P. berghei* proteins by PbGEM technology, 3xHA and mScarlet-I tagging constructs were generated using phage recombineering in *E. coli* TSA strain with PlasmoGEM vectors (http://plasmogem.sanger.ac.uk/) using sequential recombineering and gateway steps as described previously (Pfander et al., 2011). For genes SKA2 and NUF2, the Zeocin-resistance/Phe-sensitivity cassette was introduced using oligonucleotides *goi-*recR1 x *goi-*recR2 for 3xHA tagging and *goi* mSc-F x *goi* mSc-R for mSc tagging vectors. Insertion of the GW cassette following gateway reaction was confirmed using primer pairs GW1 x *goi-*QCR1 and GW2 x *goi-*QCR2.

For C-terminal tagging of *P. berghei* proteins by pCP, constructs were newly derived from pOB277 (Patzewitz et al., 2013) in order to target endogenous loci by allele replacement instead of insertion. Briefly a 588 bp fragment encompassing the coding sequence of 3xHA and DHFR flanked by KpnI and EcoRI was amplified from PbGEM plasmid GW-R6K-3xHA using primers MB1033 and MB1034 and replaced the corresponding fragment in pOB277 to generate pCP-3xHA. The coding sequence for mNG flanked by AvrII and SacII sites was purchased from GeneArt and inserted upstream to 3xHA to generate pCP-mNG-3xHA. Sequences comprising ∼500500 bp from the C-terminus of the coding sequence and ∼500 bp from the immediate 3’ UTR for genes SKA1, SKA3 and AKiTs 1 - 9 were cloned into KpnI and AvrII sites upstream to the mNG coding sequence, along with a NotI linearisation site between the targeting sequences.

For C-terminal tagging of *T. gondii*, constructs were generated by KOD PCR as described in (Brown et al. 2018). Genomic DNA extractions were performed with the Wizard SV genomic DNA purification kit (Promega). PCRs to generate specific gRNAs were performed with Q5 polymerase (New England Biolabs) while PCRs to generate specific knock-in constructs (mAID fusions and epitope tagging) were performed with KOD polymerase (Novagen). Specific gRNA were generated using the Q5 site-directed mutagenesis kit (New England Biolabs) on the pSAG1::Cas9-U6::sgUPRT vector, as described previously (Shen et al. 2014).

### Plasmodium berghei maintenance and transfection

*P. berghei* ANKA strain-derived clones 2.34 (Billker et al., 2004) together with derived transgenic lines were grown and maintained in CD1 outbred mice. Six to ten weeks-old mice were obtained from Charles River laboratories and females were used for all experiments. Mice were specific pathogen free (including *Mycoplasma pulmonis*) and subjected to regular pathogen monitoring by sentinel screening.

They were housed in individually ventilated cages furnished with a cardboard mouse house and Nestlet, maintained at 21 ± 2°C under a 12 hr light/dark cycle and given commercially prepared autoclaved dry rodent diet and water ad libitum. The parasitaemia of infected animals was determined by microscopy of methanol-fixed and Giemsa-stained thin blood smears. For gametocyte production, parasites were grown in mice that had been phenyl hydrazine treated three days before infection. Exflagellation was induced in exflagellation medium (RPMI 1640 containing 25 mM HEPES, 4 mM sodium bicarbonate, 5% fetal calf serum (FCS), 100 mM xanthurenic acid, pH 7.8). For gametocyte purification, parasites were harvested in suspended animation medium (SA; RPMI 1640 containing 25 mM HEPES, 5% FCS, 4 mM sodium bicarbonate, pH 7.20) and separated from uninfected erythrocytes on a Histodenz/Nycodenz cushion made from 48% of a Histodenz/Nycodenz stock (27.6% [w/v] Histodenz/Nycodenz [Sigma/ Alere Technologies] in 5.0 mM TrisHCl, 3.0 mM KCl, 0.3 mM EDTA, pH 7.20) and 52% SA, final pH 7.2. Gametocytes were harvested from the interface.

Schizonts for transfection were purified from overnight *in vitro* culture on a Histodenz cushion made from 55% of the Histodenz/Nycodenz stock and 45% PBS. Parasites were harvested from the interface and collected by centrifugation at 500 g for 3 min, resuspended in 25 mL Amaxa Basic Parasite Nucleofector solution (Lonza) and added to 10 µg DNA dissolved in 10 µl H2O. Cells were electroporated using the FI- 115 program of the Amaxa Nucleofector 4D. Transfected parasites were resuspended in 200 ml fresh RBCs and injected intraperitoneally into mice. Parasite selection with 0.07 mg/mL pyrimethamine (Sigma) in the drinking water (pH ∼4.5) was initiated one day after infection.

### Toxoplasma gondii maintenance and transfection

*Toxoplasma gondii* tachyzoites were grown in human foreskin fibroblasts (HFFs, American Type Culture Collection-CRL 1634) maintained in Dulbecco’s Modified Eagle’s Medium (DMEM, Gibco) supplemented with 5% fetal calf serum (FCS), 2 mM glutamine and 25 μg/ml gentamicin. Absence of *Mycoplasma* contamination was checked regularly by immunofluorescence. All mAiD fusion strains were generated in a Tir1 expressing cell line and depletion of protein achieved by incubation with 500 µM of indole-3-acetic acid (IAA) as described in (Brown et al. 2017).

Freshly egressed tachyzoites were transfected by electroporation as previously described (Soldati-Favre & Boothroyd 1993). For each transfection, 40 µg of specific gRNA was used to target the 3’UTR of the gene of interest. Mycophenolic acid (25 mg/mL) and xanthine (50 mg/mL) or pyrimethamine (1 µg/ml) were employed to select resistant parasites carrying the HXGPRT and DHFR cassette, respectively.

For assessment of protein depletion by plaque assays, HFFs were infected with fresh parasites and grown for 7 days before fixation with PFA/GA. After fixation, HFFs were washed with PBS and the host cells monolayer was stained with crystal violet.

For morphological analysis of cells depleted for kinetochore components, parasites were processed for immunofluorescence as described above and stained for tubulin, Centrin1 and DNA stains (Table S5). The mitotic index of cells was calculated in both induced and non-induced cells.

### Immunoblotting

Immunoblotting was used to confirm expression of tagged proteins. Actively dividing cells were washed in PBS and resuspended at between 1-5 × 10^5^ cells μl^-1^. Lysis was in Laemmli buffer (2% w/v SDS, 0.4 M 2-mercaptoethanol, 10% glycerol, 50 mM Tris-HCl pH 7.2) at 95°C for 5 min. Lysates containing around 5 × 10^6^ cells were separated on 4-20% polyacrylamide gels (Invitrogen) in running buffer (25 mM Tris, 250 mM glycine and 0.1% w/v SDS). Proteins were electrophoretically transferred to 0.45 μm pore-size nitrocellulose membrane at 1.6V cm^-1^ for 14 h in transfer buffer (25 mM Tris, 192 mM glycine, 0.02% w/v SDS and 10-20% methanol). Membranes were blocked and in 5% w/v milk powder in TBS-T (20 mM Tris-HCl pH 7.5, 150 mM NaCl, 0.05% Tween-20) for 1 h. Membranes were incubated in primary antibody (Table S5) in 1% w/v milk in TBS-T for 1 h and washed in TBS-T. Detection was by secondary peroxidase-conjugated antibody (Table S5), in 1% w/v milk in TBS-T for 1 h. Membranes were washed and detected by chemiluminescence with Western Lightning ECL (PerkinElmer) and exposure to photographic film.

### Protein localization

For localization of tagged proteins in *P. berghei* by native fluorescence, cells from different proliferative stages during the parasite lifecycle were mounted in exflagellation medium (RPMI 1640 containing 25 mM HEPES, 4 mM sodium bicarbonate, 5% fetal calf serum (FCS), 100 mM xanthurenic acid, pH 7.8), and imaged using an inverted Zeiss Axio Observer Z1 microscope fitted with an Axiocam 506 mono 14 bit camera and Plan Apochromat 63x / 1.4 Oil DIC III objective.

For immunofluorescence of *T. gondii*, parasites were inoculated on an HFF monolayer with coverslips in 24-well plates, grown for 16–30 hours depending on the experiment, and fixed with cold methanol (- 20°C) for 7 min. Coverslips were then washed with PBS and blocked for 20 min in PBS-BSA 5%. Primary and secondary antibodies (Table S5) were then incubated sequentially for 1 hour each in PBS-BSA 2%. Three washes of 5 minutes each were performed between primary and secondary antibodies incubations, using PBS. Coverslips were mounted in VECTASHIELD® Antifade Mounting Medium with DAPI and imaged using the inverted Zeiss Axio Observer, as described above.

For ultrastructure expansion microscopy (U-ExM), cells were sedimented on poly-D-lysine (A-003-E, Sigma) coverslips (150 μL/coverslip) during 10 minutes at room temperature (RT) and fixed in −20°C methanol during 7 minutes and prepared for U-ExM as previously published (Bertiaux et al., 2021). Briefly, coverslips were incubated for 5 hours in 2X 1.4% AA/ 2% FA mix at 37°C prior gelation in APS/Temed/Monomer solution (19% Sodium Acrylate; 10% AA; 0,1% BIS-AA in PBS 10X) during 1 hour at 37°C. Gels were denatured at 95°C for 1 hour. Gels were incubated in ddH2O overnight for expansion. The following day, gels were washed in PBS before incubation with primary antibodies (Table S5) for 3 hours at 37°C. Gels were washed in PBS-Tween 0.1% prior incubation with secondary antibodies (Table S5) for 3 hours at 37°C. Gels were washed in PBS-Tween 0.1%. Gels were incubated in ddH2O for a second round of expansion before imaging. For NHS-ester staining, directly after antibody stainings gels were incubated in NHS-Ester (Thermo Fisher Scientific catalog number: 46402) 10 μg/mL in PBS for 1 hour and 30 minutes at RT on a rocking platform and washed 3x in PBS before overnight expansion. Confocal microscopy was performed on a Inverted Leica DMi8 with a HC PL Apo 100x/ 1.40 Oil immersion objective fitted with a Leica DFC7000T camera.

All images of fluorescent proteins were captured at RT with equal exposure settings. Images for level comparison were processed with the same alterations to minimum and maximum display levels. For analysis of relative positions of kinetochore components, peak locations of tagged proteins of focus centroids in each channel were calculated. Analysis was performed in Fiji (Schindelin et al., 2012) and the statistical programming package R (http://www.r-project.org). Distances were always measured from dual staining experiments. The expansion factor of each gel was after applied to the value to obtain the real distance.

### Immunopurification

For the purification of kinetochore complexes, gametocytes were purified from 5-10 ml *P. berghei* infected mouse blood. Cells were washed twice in HKMEG (150 mM KCl, 150 mM glucose, 25 mM HEPES, 4 mM MgCl2, 1 mM EGTA, pH7.8) containing 100 x protease inhibitor cocktail and 20 μM MG132. Sequential chemical cross-linking IP followed as described (D’Archivio & Wickstead, 2017). Briefly, cells were treated with 0, 0.1, or 1% formaldehyde in HKMEG for 10 min, quenched with 10 ml of 1 M glycine, and lysed in HKMEG containing 1% (vol/vol) Nonidet P-40, 0.5% sodium deoxycholate, 0.1% SDS, 1mM dithiothreitol (DTT), 100 x protease inhibitor cocktail and 20 μM MG132. Lysate was sonicated in an Ultrasonic water bath for 20 min applied for 50% of the cycle and cleared by centrifugation at >20,000 g for 30 min. The soluble fraction was then allowed to bind to affinity-purified rat anti-HA antibody (Sigma) attached to paramagnetic beads (Dynabeads protein G; Invitrogen) for 4 hours. Beads were washed in HKMEG containing 0.1% (vol/vol) Nonidet P-40, 0.5mM dithiothreitol. Beads were re-suspended in 100 µl of 6 M urea in 50 mM ammonium bicarbonate (AB). 2 µl of 50 mM dithioerythritol (DTE) were added and the reduction was carried out at 37°C for 1 hr. Alkylation was performed by adding 2 µl of 400 mM iodoacetamide for 1 hr at room temperature in the dark. Urea was reduced to 1 M by addition of 500 ml AB and overnight digestion was performed at 37°C with 5 ml of freshly prepared 0.2 mg/ml trypsin (Promega) in AB. Supernatants were collected and completely dried under speed-vacuum. Samples were then desalted with a C18 microspin column (Harvard Apparatus) according to manufacturer’s instructions, completely dried under speed-vacuum and stored at -20°C.

### Mass spectrometry

Protein identification followed as described (Balestra et al., 2020). Samples were diluted in 20 µl loading buffer (5% acetonitrile, 0.1% formic acid [FA]) and 2 µl were injected onto the column. LC-ESI-MS/MS was performed either on a Q-Exactive Plus Hybrid Quadrupole-Orbitrap Mass Spectrometer (Thermo Fisher Scientific) equipped with an Easy nLC 1000 liquid chromatography system (Thermo Fisher Scientific) or an Orbitrap Fusion Lumos Tribrid mass Spectrometer (Thermo Fisher Scientific) equipped with an Easy nLC 1200 liquid chromatography system (Thermo Fisher Scientific). Peptides were trapped on an Acclaim pepmap100, 3 µm C18, 75 µm x 20 mm nano trap-column (Thermo Fisher Scientific) and separated on a 75 µm x 250 mm (Q-Exactive) or 500 mm (Orbitrap Fusion Lumos), 2 µm C18, 100 Å Easy-Spray column (Thermo Fisher Scientific). The analytical separation used a gradient of H 2O/0.1% FA (solvent A) and CH3CN/0.1% FA (solvent B). The gradient was run as follows: 0 to 5 min 95% A and 5% B, then to 65% A and 35% B for 60 min, then to 10% A and 90% B for 10 min and finally for 15 min at 10% A and 90% B. Flow rate was 250 nL/min for a total run time of 90 min. Data-dependent analysis (DDA) was performed on the Q-Exactive Plus with MS1 full scan at a resolution of 70,000 Full width at half maximum (FWHM) followed by MS2 scans on up to 15 selected precursors. MS1 was performed with an AGC target of 3 x 10^6^, a maximum injection time of 100 ms and a scan range from 400 to 2000 m/z. MS2 was performed at a resolution of 17,500 FWHM with an automatic gain control (AGC) target at 1 x 10^5^ and a maximum injection time of 50 ms. Isolation window was set at 1.6 m/z and 27% normalised collision energy was used for higher-energy collisional dissociation (HCD). DDA was performed on the Orbitrap Fusion Lumos with MS1 full scan at a resolution of 120,000 FWHM followed by as many subsequent MS2 scans on selected precursors as possible within a 3 s maximum cycle time. MS1 was performed in the Orbitrap with an AGC target of 4 x 10^5^, a maximum injection time of 50 ms and a scan range from 400 to 2000 m/z. MS2 was performed in the Ion Trap with a rapid scan rate, an AGC target of 1 x 10^4^ and a maximum injection time of 35 ms. Isolation window was set at 1.2 m/z and 30% normalised collision energy was used for HCD.

### Label-free quantitation

Label-free quantitation followed as described (Brusini et al., 2021; D’Archivio & Wickstead, 2017) and performed on .mgf data files using the Central Proteomics Facilities Pipeline (cpfp.sourceforge.io). Data were searched with X!Tandem and OMS SA engines against a custom, nonredundant protein database of predicted protein sequences from ANKA 2.34 strain including exogenous protein sequences (fluorescent proteins, drug selection markers, and exogenous proteins expressed in the parental cells) and common contaminating peptides. Possible modification of peptides by N-terminal acetylation, phosphorylation (P), ubiquitination (Ub), carbamidomethylation (C), oxidation (M), and deamidation (N/Q) was permitted in searches. Peptide identifications were validated with PeptideProphet and ProteinProphet (Nesvizhskii et al., 2003) and lists compiled at the peptide and protein level. iProphet was used to combine search engine identifications and refine identifications and probabilities. Normalized spectral index quantitation (SINQ) was applied to the grouped metasearches to give protein-level quantitation between labelled samples and controls (Trudgian et al., 2011), and implemented by the Central Proteomics Facilities Pipeline. SINQ values are summed intensities of matched fragment ions for all spectra assigned to a peptide (identified by ProteinProphet), normalized for differences in protein loading between datasets and for individual protein length. Proteins with 1 detected peptide and an estimated false discovery rate of ≤ 1% relative to a target-decoy database were considered. A total of 780 distinguishable *Plasmodium* proteins were detected across all experiments. Mass spectrometry proteomics data have been deposited to the ProteomeXchange Consortium via the PRIDE (Perez-Riverol et al., 2019) partner repository with the dataset identifier PXD028595. Processed data are also provided in Table S2.

Enrichment and principal component analyses were performed in the statistical programming package ‘R’ (www.r-project.org). Quantitative values were analyzed as either log-transformed SINQ values (for principal component analysis) or log-transformed ratio of sample SINQ value versus control immuno- purification (enrichment analysis).

### Bioinformatic analyses

Searches were based on predicted protein datasets for 90 alveolate organisms from genomes and transcriptomes (Table S1), in addition to kinetochore protein sequences identified previously (van Hooff et al., 2017). For transcriptomes, ORFs were predicted using TransDecoder. Initial profiles for each identified apicomplexan kinetochore protein were aligned using MAFFT (Katoh et al., 2005), the L-INS-i strategy, trimmed to conserved regions with trimAl, modelled by HMMER3 (Eddy, 2009) and searched to find similar sequences in all predicted proteomes and kinetochore sequences. Hits were added iteratively and used to create new profiles until no new sequences were identified. HMMs for all apicomplexan orthologous groups were also generated and defined by OrthoFinder (Emms & Kelly, 2019). Alignments were visualized and modified using Jalview (Waterhouse et al., 2009). Profile–profile comparisons were performed using HH-suite3 (Söding, 2005). For phylogenies, trimmed alignments were used to infer maximum likelihood phylogenies using IQ-TREE2 (Minh et al., 2020). Trees were visualized in FigTree (tree.bio.ed.ac.uk/software/figtree) and Phytools (Revell, 2012), as part of the statistical programming package ‘R’ (www.r-project.org).

